# Generation and propagation of network bursts in the basal ganglia exhibit dynamic changes during early postnatal development

**DOI:** 10.1101/2022.07.26.501540

**Authors:** Sebastian Klavinskis-Whiting, Sebastian Bitzenhofer, Ileana Hanganu-Opatz, Tommas Ellender

**Affiliations:** Department of Pharmacology University of Oxford Mansfield Rd Oxford, OX13QT, United Kingdom; Department of Biomedical Sciences University of Antwerp Universiteitsplein 1 2610 Wilrijk, Belgium; Developmental Neurophysiology Institute of Neuroanatomy University Medical Center Hamburg-Eppendorf 20246 Hamburg, Germany

**Keywords:** development, network bursts, oscillations, spindle bursts, nested gamma spindle bursts, striatum, cortex, intralaminar thalamus

## Abstract

The neonatal brain is characterised by intermittent bursts of oscillatory activity interspersed by relative silence. While these bursts of activity are well characterised for many cortical areas much less is known whether and how these propagate and interact with subcortical regions. Here, early network activity was recorded using silicon probes from the developing basal ganglia, including the motor/somatosensory cortex, dorsal striatum and intralaminar thalamus, during the first two postnatal weeks in mice. Using an unsupervised detection and classification method, two main classes of bursting activity were found, consisting of spindle bursts (SB) and nested gamma spindle bursts (NGB), which were characterised by oscillatory activity at respectively ∼10 Hz and ∼30 Hz. These bursts were reliably identified across all three brain structures but differed in their structural, spectral, and developmental characteristics. Coherence and cross-correlation analyses revealed that burst events often occur synchronously across different brain regions and were mostly of a similar type, especially between cortex and striatum, which also exhibited the strongest interactions as compared to other brain regions. Interestingly, the preferred frequency for these interactions suggested a developmental shift from initial lower frequencies to higher frequencies across development. Together, these results provide the first detailed description of early network activity within the developing basal ganglia and suggests that distinct brain regions drive and coordinate burst activity at different developmental stages.

## Introduction

The developing brain is a hugely dynamical system, with marked changes in structural organization and functional connectivity occurring on both fast (seconds) and slow (days) timescales (Stiles & Jernigan, 2010). It has been demonstrated that transient bursts of neural activity, which characterize the brain during this dynamic period, are critical for many developmental processes including neuronal migration and synaptic refinement (Katz & Shatz, 1996; Heck *et al*., 2008; Bonetti & Surace, 2010; Yasuda *et al*., 2011; Bando *et al*., 2016; Minocha *et al*., 2017; Suchkov *et al*., 2018; Yasuda *et al*., 2021). Disruption of these early patterns of activity can lead to aberrant neuronal circuit development and lasting cognitive impairments in animal models (Hartung *et al*., 2016b; Chini & Hanganu-Opatz, 2020; Chini *et al*., 2020; Bitzenhofer *et al*., 2021) and likely contribute also to the emergence of a variety of neurodevelopmental and neurological disorders including schizophrenia, autism and epilepsy, amongst others (Graybiel & Rauch, 2000; Del Campo *et al*., 2011; Langen *et al*., 2011; McNaught & Mink, 2011; Shepherd, 2013; Albin, 2018; Chini & Hanganu-Opatz, 2020; Molnar *et al*., 2020; Iannone & De Marco Garcia, 2021; Luhmann *et al*., 2022). Many studies of these patterns of activity have been performed in cortical structures (Hanganu *et al*., 2006; Brockmann *et al*., 2011; An *et al*., 2014; Cichon *et al*., 2014; Shen & Colonnese, 2016) or related afferent structures such as thalamus (Weliky & Katz, 1999; Minlebaev *et al*., 2011; Yang *et al*., 2013). In these brain regions various activity patterns have been described, which are typically classified based on their distinct properties (e.g. amplitude, duration) and spectral structure, and can either be spontaneously generated (Hanganu *et al*., 2006; Yang *et al*., 2013; Luhmann *et al*., 2016; Martini *et al*., 2021) or evoked by external sensory inputs (Khazipov *et al*., 2004; Milh *et al*., 2007; Yang *et al*., 2013; Gerasimova *et al*., 2014). These transient bursts of activity largely disappear at later stages of development and are replaced by longer duration oscillatory activity whose spectral components have themselves been related to various physiological and cognitive functions (Boraud *et al*., 2005; Colgin, 2013; Khazipov *et al*., 2013).

How these transient patterns of early bursting activity impact or interact with many other parts of the developing brain, including the basal ganglia, is largely unknown. The basal ganglia consist of a group of subcortical brain nuclei essential for the control of movement, as well as a variety of other functions in adulthood (Graybiel *et al*., 1994; Grillner *et al*., 2005; Yin & Knowlton, 2006). It is highly likely that early patterns of activity propagate through the basal ganglia as they can travel widely and coordinate patterns of activity amongst widely separated brain regions (Ackman *et al*., 2012; Hartung *et al*., 2016a; Ahlbeck *et al*., 2018). The striatum is the principal input nucleus of the basal ganglia which receives its main excitatory synaptic inputs from the cortex and thalamus (Buchwald *et al*., 1973; Smith *et al*., 2004; Kreitzer, 2009; Doig *et al*., 2010; Ellender *et al*., 2013; Smith *et al*., 2014; Hunnicutt *et al*., 2016). While the cortico-striatal inputs come from widespread, and extensive cortical areas (Oh *et al*., 2014; Hunnicutt *et al*., 2016), the main thalamic inputs to the striatum are thought to arrive predominantly from the intralaminar nuclei (Macchi *et al*., 1984), which in rodents can be divided in the rostral central lateral (CL) and the caudal parafascicular (Pf) nucleus (Berendse & Groenewegen, 1990; Smith *et al*., 2009). Input from primary sensory thalamic nuclei input is thought to be minimal to non-existent to striatum at least in adulthood (Alloway *et al*., 2017; Ponvert & Jaramillo, 2019). Cortical activity can also indirectly impact striatum through the Pf thalamic nucleus as it receives strong direct innervation from the cortex as part of topographically segregated thalamocortical loops (Mandelbaum *et al*., 2019). These excitatory synaptic inputs innervate both the striatal spiny projection neurons (SPNs) and interneurons (Sadikot *et al*., 1992; Castle *et al*., 2005; Lacey *et al*., 2007; Doig *et al*., 2010; Ellender *et al*., 2013; Smith *et al*., 2014) and become functional in the first postnatal weeks (Tepper *et al*., 1998; Nakamura *et al*., 2005; Dehorter *et al*., 2011; Peixoto *et al*., 2016; Krajeski *et al*., 2019). Appropriate levels of neural activity in the basal ganglia are necessary for the proper development of synapses and circuits in the early postnatal striatum (Kozorovitskiy *et al*., 2012; Peixoto *et al*., 2016; Peixoto *et al*., 2019) as well as the survival of striatal neurons (Sreenivasan *et al*., 2022). Indeed, aberrant levels of activity during development can cause permanent striatal dysfunction (Mowery *et al*., 2017; Vicente *et al*., 2020). Despite these clear roles for early neural activity, we still know very little about the diversity of activity patterns that dominate the basal ganglia during early postnatal development, including their initiation and propagation and possible interactions amongst basal ganglia nuclei.

Here, we use multi-site *in vivo* silicon probes to record early network activity in young mice during the first postnatal weeks (postnatal day (P) 5 to P15) from three key nodes in the basal ganglia network, including the motor/somatosensory cortex, the intralaminar thalamic nuclei and the dorsal striatum. We focussed on these three different regions to respectively capture the two main excitatory afferent structures to the basal ganglia, as well as the main input nucleus of the basal ganglia. We applied an unbiased detection and classification method to these patterns of activity and explored to what extent these activity patterns propagate and interact in the distinct brain regions and developmental time periods. We find that patterns of brief bursting activity in the developing basal ganglia are dominated by spindle (∼10Hz) and beta-low gamma (∼15-30Hz) frequency bands, corresponding to spindle bursts and nested gamma spindle bursts respectively, which were found in all three key nodes. These exhibited subtly different characteristics and developmental profiles, and interestingly our analysis suggests these underpinned distinct directional interactions amongst brain regions during early postnatal development.

## Methods & Materials

### Animals

All experiments were carried out on C57Bl/6 wildtype mice of both sexes with *ad libitum* access to food and water. Experiments were designed to use litter mates for extended developmental ranges within single experiments, as to control for effects of litter sizes and maternal care factors that could affect the degree of neuronal and circuit maturity. All mice were bred, IVC housed in a temperature-controlled animal facility (normal 12:12 h light/dark cycles) and used in accordance with the UK Animals (Scientific Procedures) Act (1986).

### Surgery and acute extracellular recording conditions

All experiments were conducted in accordance with the national laws and the UK guidelines for the use of animals in research and approved by the local ethical committee. Multi-site extracellular silicon probe recordings were performed unilaterally in cortex and striatum (0.5 mm posterior to bregma and 3.5 mm from the midline at a 30° angle) and thalamus (1.0-1.5 mm anterior to bregma and 0.7 mm from the midline) of postnatal day 5 (P5) to P15 mice using experimental protocols as described previously (Hanganu *et al*., 2006; Brockmann *et al*., 2011). Briefly, urethane was injected intraperitoneally (1 mg/g body weight) prior to the surgery and under isoflurane anaesthesia a plastic bar head mount was fixed with dental cement to the nasal and occipital bones. A craniotomy was performed above the recording locations, taking care not to pierce the dura mater, by drilling a hole of ∼0.5-1 mm in diameter. The head was fixed into the stereotaxic apparatus (Stoelting) using the plastic bar head mount. During recordings, the body of the animals was surrounded by cotton and kept at a constant temperature of 37° C using a heating blanket. Multi-site electrodes (NeuroNexus, MI, USA) were inserted into the brain (A16, 5 mm length, cortex/striatum: 100 μm spacing and thalamus: 50 μm spacing). Silver wires were inserted into the cerebellum and served as ground and reference. After 30 min recovery the recordings were started from silicon probes to obtain simultaneous recordings of field potential (FP) and multiple-unit activity (MUA) at different depths and locations. The electrodes were labelled with DiIC18(7) (1,1’-Dioctadecyl-3,3,3’,3’-Tetramethylindotricarbocyanine Iodide) (DiR) (Invitrogen) to enable post-mortem reconstruction of the electrode tracks in histological sections.

### Data acquisition

Recordings lasted between 30 - 60 minutes. Data were acquired at a sampling rate of 30 kS/s using the Open Ephys acquisition system (Siegle *et al*., 2017). Silicon probes were connected to electrode adapter boards to 16-channel recording headstages (Intan Technologies) which were each connected to the Open Ephys acquisition board which was connected to a Dell Latitude 7370 laptop.

### Data analysis

Data were analyzed offline using custom written scripts in Igor Pro (Wavemetrics, RRID:SCR_000325) and Python. Analysis code is accessible at https://github.com/sebbkw/networkbursts2022. Data were filtered offline at 1-100 Hz for local field potential (LFP) analyses or 400-4000 Hz for multi-unit activity (MUA) analyses using a 3rd order Butterworth filter. From each recording session, the data channel for analysis was selected out of three channels which best coresponded to the required anatomical location (i.e. cortex, dorsal striatum and intralaminar thalamus) and exhibited the lowest amount of noise by visual inspection. In addition, in three recording sessions the presence of electrical noise artefacts was deemed too large, and these were not used for subsequent analysis.

#### Spike detection

Spikes were detected by applying a threshold set at 5 standard deviations below baseline MUA (Muthmann *et al*., 2015). Candidate spike waveforms were extracted by selecting the 30 time-samples (1 ms) surrounding the local minimum over each set of consecutive time points crossing the threshold. Spikes were rejected if the maximum point did not exceed 0 µV.

#### Burst detection

Two burst detection methods — envelope thresholding and root mean square (RMS) thresholding — were initially trialled (see Supplemental Figure 1). For both methods, the LFP signal was first band-pass filtered between 4-100 Hz to remove low-frequency artefacts which could bias the detection of bursts. Since no ground truth is available (i.e., there is no objective definition for which time periods constitute bursting events), the use of two independent methods provided an additional means of validating burst detection by considering whether the two methods produced comparable results. Overall, overlap was good for striatum (mean overlap: 84%, SD: 11%) and thalamus (mean overlap: 82%, SD: 14%), indicating that both successfully identified burst events with some method-specific variability. Ultimately, the RMS thresholding method was chosen for all subsequent analyses as it has been better validated in the literature and deemed more parsimonious insofar as it does not require any per-recording parameters to be adjusted by the experimenter. At around P12, a developmental switch was observed whereby bursting events were no longer discernible from continuous oscillatory activity. Accordingly, all subsequent analyses for bursting events were only performed using data from animals aged P5-12. The RMS thresholding method was adapted from Cichon and colleagues (Cichon *et al*., 2014). In brief, RMS values were first computed for each 200 ms window of the LFP signal before being binned to produce a distribution of RMS values. The RMS threshold was determined by fitting a Gaussian function to this RMS distribution, taking the threshold as *µ*+3*σ* where *µ* is the mean and *σ* is the standard deviation of the fitted Gaussian. Burst events were defined as those 200 ms time periods where the RMS exceeded the RMS threshold. At older ages (i.e. P>10) where the proportion of bursting events relative to baseline periods increased, a naive curve-fitting procedure tended to overestimate the optimal threshold such that only the peaks of bursting events were detected. Therefore, and unlike for Cichon *et al*. (2014), the Gaussian mean was limited to a maximum RMS value of 50 µV during curve fitting. All consecutive 200 ms periods which exceeded the threshold were combined and defined as a single burst event. Several criteria were applied to further ensure the quality of detected bursts and to minimise the occurrence of false positives (i.e. periods of silence identified as burst events). Firstly, putative burst events were rejected if they were below 0.2 s or above 20 s in duration. Although exceedingly long (40-80 s) spindle-like bursts have been described in certain parts of cortex (Yang *et al*., 2013) no such bursts were observed in our recordings by visual inspection. Moreover, as most bursting activity in cortex in general has been shown to be relatively transient in the order of 1-10 s (Minlebaev *et al*., 2011; Khazipov *et al*., 2013; Gerasimova *et al*., 2014; Suchkov *et al*., 2018) we excluded bursts exceeding 20 s from analysis. The total percentage of rejected events was 1.3% for cortex, 2% for striatum and 1.3% for thalamus. Secondly, bursting events were required to have at least 5 peaks above the mean RMS for that burst event to exclude non-specific increases in LFP amplitude not accompanied by oscillatory activity. Thirdly, motion artefacts were accounted for by rejecting all events with a mean RMS value exceeding 1000 µV.

#### Unsupervised burst classification

Burst events were classified using a combined principal component analysis (PCA) and clustering approach. PCA was used as a feature extraction method to identify latent variables in the feature space which could then be used to cluster bursting events into different groupings. This data-driven approach was both unsupervised and unbiased as compared to manual classification methods, since bursts were classified using those axes that explained maximal variance in the data rather than relying on predefined criteria. Each burst event was used to produce a feature vector on which the PCA procedure was performed. Nine features were included for the analysis: duration, negative peak, maximum RMS, flatness, maximum slope, inter-trough interval (ITI), relative theta-alpha power, relative beta-low gamma power and spike rate. These distinct features were selected to capture a variety of aspects of each bursting event including their structural and spectral characteristics as validated in previous work (Cichon *et al*., 2014; Hartung *et al*., 2016a; Hartung *et al*., 2016b). For all features other than relative theta-alpha power and relative beta-low gamma power, burst events were band-pass filtered at 4-100 Hz. Duration was computed as the difference between the start and end times of the detected burst event. The negative peak was defined as the minimum deflection in the LFP signal. Maximum RMS was defined as the maximum RMS values computed across 200 ms chunked time windows. Flatness was defined as min(RMS)*/*max(RMS) out of all 200 ms period RMS values across the burst event. Slope was taken as the instantaneous difference in voltage between consecutive time points, for which the burst event was downsampled to 500 S/s to ensure that inter-sample time points were large enough to produce sufficient variation in slope across burst events. The ITI was defined as the mean time interval between all troughs in the LFP signal, where local minima were included as troughs only if their prominence exceeded half the RMS of the burst event. Spectral measures were taken as the power in the theta-alpha (4-16 Hz) and beta-low gamma (16-40 Hz) bands relative to the total power of the normalized power spectra in the frequency range 1-50 Hz. Spike rate was defined as the number of spike events occurring per second over the course of the burst event. Prior to running the PCA, all features were normalized. Bursts were then clustered using the first three components extracted by the PCA. Clustering was performed using the fuzzy c-means clustering algorithm, where — in contrast to hard clustering algorithms — each point is assigned a relative probability for its inclusion to each cluster (Ross, 2010). By virtue of this ‘fuzzy’ approach, clustering could make use of a threshold whereby points of low confidence for inclusion to any cluster were labelled as unclassified. For clustering in this study, burst events were labelled as unclassified if the maximum coefficient did not exceed 0.6, indicating less than 60% probability of the burst belonging to either of the two candidate clusters. The clustering threshold was set as a compromise between ensuring that low-confidence bursts were not included in either cluster and that excessive burst events were not unduly discarded from further analysis. The optimal number of clusters was determined by measuring the fuzzy partition coefficient (FPC) as a function of cluster number, where the FPC denotes the quality of the resulting partition with an optimal value of 1. Excluding the trivial case where *n* = 1, the optimal number of clusters was determined as *n* = 2 for both striatal (FPC = 0.75) and thalamic (FPC = 0.73) burst events (Supplemental Figure 3).

#### Time-frequency analysis

LFP power spectra were estimated using the multi-taper method via the ‘pmtm’ function in the Spectrum toolbox (Cokelaer & Hasch, 2017). For all power spectral density (PSD) analyses, 1 s windows were advanced by 0.1 s with a half-bandwidth parameter of 3 using the first 5 Slepian sequences. To account for non-specific spectral properties during bursting events, all PSDs were normalized (*P/P*_0_) according to the PSD measured during baseline non-bursting time periods (Cichon *et al*., 2014; Gretenkord *et al*., 2019). The baseline PSD was calculated as the mean PSD across all periods in each recording not included as bursting events, which exceeded 1 s in duration. Spectrograms were produced purely for display purposes using the SciPy ‘spectrogram’ function in the signal toolbox, using a time window of 0.5 s with a 99% overlap across segments.

#### Coherence analyses

Bursts were considered to be co-occurring if their onset overlapped by less than 0.5 s. The frequency of co-occurring burst types (SB/SB, NGB/NGB, NGB/SB) was normalized by the total number of bursting events in each class to control for their absolute incidence. Cross-spectral coherence between co-occurring burst events was then computed using Welch’s method using the ‘coherence’ function in the SciPy signal toolbox (Virtanen *et al*., 2020). Cross-spectral coherence was computed with a time window of 0.5 s and 0 s of overlap across segments. Significance was determined by shuffling all pairs of burst events 1000 times and computing the mean coherence at each iteration to produce a null distribution (see Supplemental Figure 8). The significance threshold was calculated by computing the 95^th^ percentile of the resulting null distribution. Co-occurring events with a mean cross-spectral coherence exceeding 0.8 were rejected, as these events of high synchrony are likely to have resulted from artefacts such as movement (Hartung *et al*., 2016a).

Imaginary coherence was computed as the absolute of the imaginary component of the normalized cross-spectrum using Welch’s with a time window of 0.5 s and 0 s of overlap across segments (Gretenkord et al., 2019). Significance was determined by shuffling all pairs of burst events 1000 times and computing the mean coherence at each iteration to produce a null distribution.

Spike-field coherence was computed as previously described (Soteropoulos & Baker, 2006). Spike trains were binned in 2 ms time-segments to produce a continuous waveform, with LFP signals (4-80 Hz bandpass filtered) similarly down sampled to match the spike-train sampling rates. Cross-spectral coherence between these two signals was then computed as the real component of the normalized cross-spectrum as described above. The significance of synchrony was assessed by constructing a null distribution across 1000 iterations where spike timing were randomly shuffled.

#### Cross-correlation analyses

Causal interactions between brain regions were assessed by means of a cross-correlation analysis (Adhikari *et al*., 2010; Hartung *et al*., 2016a). First, LFP signals were band-pass filtered around the frequencies of interest – here, the 4-16 Hz and 16-40 Hz frequency windows around which spectral power was greatest for SB and NGB bursts, respectively. The LFP signal was then pre-whitened to remove autocorrelations which can produce spurious correlations between input signals (El-Gohary & McNames, 2007). More specifically, pre-whitening was achieved by fitting an ARIMA model to the LFP signal before computing the normalized cross-correlation on the resulting residuals of the fitted model (Merchant *et al*., 2014). The lag was then taken as the time lag between +50 and −50 ms, which produced the maximal cross-correlation. For cortical-striatal bursts, a positive lag indicates the striatum leading the cortex and a negative lag the cortex leading the striatum. For thalamic-striatal bursts, a positive lag indicates the striatum leading the thalamus and a negative lag the thalamus leading the striatum. Finally, for cortical-thalamic bursts, a positive lag indicates the thalamus leading the cortex and a negative lag the cortex leading the thalamus. Shorter lags of less than 20 ms were considered putative monosynaptic interactions, while longer lags were indicative of possible polysynaptic interactions (Hartung *et al*., 2016a).

#### Spike train cross-correlation/lag analysis

Spike times from the co-occurring burst events were first extracted from the filtered MUA signal as described above, after which spike trains were convolved with a Gaussian kernel with a standard deviation of 2 ms to produce a continuous signal. The normalized cross-correlations of these two signals were then computed and the peak lag taken between +/- 10 ms. Significance was assessed using a one-sample *t*-test across the peak lag from each burst pair.

### Histological analyses

Following recordings, the mice were culled, and brains immediately transferred to ice-cold 4% paraformaldehyde in 0.1 M PBS and fixed for 3 or more days. Brains were then sectioned on a Leica VTS1000 vibratome and coronal sections (40 µm) collected. All sections were mounted in Vectashield (Vector Laboratories, Cat. H-1000) and DiR recording tracts were immediately analysed. Images were captured with a Leica epifluorescence microscope using HCImage software (Hamamatsu). Images were processed and analysed in ImageJ and Adobe Photoshop and Illustrator.

### Statistics

Statistics were computed using data binned into two-day age groups (namely P5-6, 7-8, 9-10, 11-12). Recordings were made from a total of 23 animals and in most, but not all, cases consisted of simultaneous recordings from cortex/striatum and thalamus for (P5-6; n=4 mice cortex/striatum and n=3 mice also including thalamus, P7-8; n=3 and n=2, P9-10; n=8 and n=5, P11-12; both n=4, P13-15; both n =4. Age-dependent effects were assessed by means of two-way ANOVA with factors of age group and either burst-type (SB and NGB) or brain region (cortex, striatum, and thalamus). To assess the dependence of burst features on brain area, we compared the same burst type across different brain areas using a non-parametric approach to multivariate analysis of variance (Anderson, 2001). Although formally equivalent to a *t*-test where the number of groups *K* = 2, this multivariate approach avoided the need for many multiple comparisons across groups and features. As for the univariate case, the F statistic was computed as:

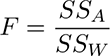

*SS_A_* is the ‘among’ or between group sum-of-squares, defined as:

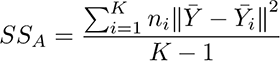

where *K* is the total number of groups (here, 2), *n_i_* is the number of observations in group *i* and ‖*Y̅*-*Y̅_i_*‖ is the Euclidean distance between the overall mean feature vector *Y̅* and the group mean feature vector *Y̅_i_*. *SS_W_* is the within group sum-of-squares, defined as:

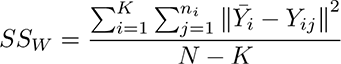

Where ‖*Y̅_i_*-*Y_ij_*‖ is the Euclidean distance between individual feature vector *Y_ij_* and group mean feature vector *Y̅_i_* and *N* is the total number of feature vectors across all groups. To assess the significance of the F-statistic, groups were shuffled 10,000 times, computing the test statistic each time to produce a null distribution against which the significance of the true F-statistic could be determined. Further statistical details of experiments can be found in the respective Results sections and Figure legends. For all statistical tests, the significance level was set at α = 0.05, with p-values corrected for multiple comparisons using Bonferroni correction where relevant. All error bars are given as standard error of the mean (SEM) except where stated otherwise. Absolute p-values are given for all values except for permutation tests where the lower bound was based on the number of permutations (e.g. p<0.0001 for 10000 permutations). Significance is indicated in Figures as * p<0.05, ** p<0.01, *** p<0.001.

## Results

### Multi-channel recordings of early neural activity in the developing basal ganglia

The basal ganglia are a complex interconnected network of subcortical nuclei of which the striatum is the main input nucleus receiving excitatory synaptic inputs from extended regions of cortex as well as diverse thalamic nuclei (Hunnicutt *et al*., 2016). How during early development activity patterns propagate and interact with each other in the different basal ganglia regions is largely unknown. Here, recordings of neural activity were made using two 16-channel silicon probes placed in the motor/somatosensory cortex, dorsal striatum as well as the intralaminar thalamus, including both the central lateral (CL) and parafascicular (Pf) nuclei, in young postnatal mice ranging in age from postnatal day (P)5 to P15 (Figure 1A). Recordings were made from a total of 23 animals under urethane anaesthesia, which is thought to minimally interfere with brain activity and its spectral characteristics (Chini *et al*., 2019). These revealed extended periods of quiescence interspersed with oscillatory events as detected from subsets of channels reflecting neuronal activity within specific brain regions (Figure 1B). These recordings were analysed using unbiased burst detection algorithms (see **Methods**, Supplemental Figure 1 and Cichon et *al.*, 2014). Initial analysis of recordings suggested that neural activity, as assessed at the level of local field potentials (LFP) and multi-unit activity (MUA), exhibited gradual changes through postnatal development. At P5, LFP activity was characterised by mostly electrical silence, interspersed with small deflections likely reflecting the first beginnings of synchronised neural activity (Figure 1C). At later stages, bursting events became more apparent as larger transient increases in synchronous activity in the LFP signal interspersed by periods of silence. Finally, by the start of the third postnatal week the periods of silence became more infrequent, and the brain state became predominantly active, indicating a more mature state akin to that found also in the adult brain (Figure 1C, D). The overall developmental trajectory appeared similar across cortex, striatum, and thalamus, indicating a gradual progression from a relatively quiet brain state towards one that generates intermittent bursts of activity before the onset of a continuous active state. Indeed, the incidence of bursts in all three brain regions increased significantly across development (F(4, 42) = 12.1, p = 0.000001) before declining at later stages of development (>P12) when all areas began to exhibit continuous oscillatory activity (Figure 1C, D). Initial analysis of data filtered to reveal spikes (0.4-4 kHz) demonstrated an overall increase in spiking activity in all three brain regions (F(4,45) = 8.83, p = 0.000024, Figure 1C, E). The increase in activity within these different brain regions was not only evident in overall increases in spike frequency and bursting incidence but was also reflected in other measured parameters such as burst amplitude and the proportion of recording time occupied by burst events (Supplemental Figure 2). All further analysis was performed on recordings up to P12 and was focussed on the intermittent bursting activity only.

**Figure 1:**
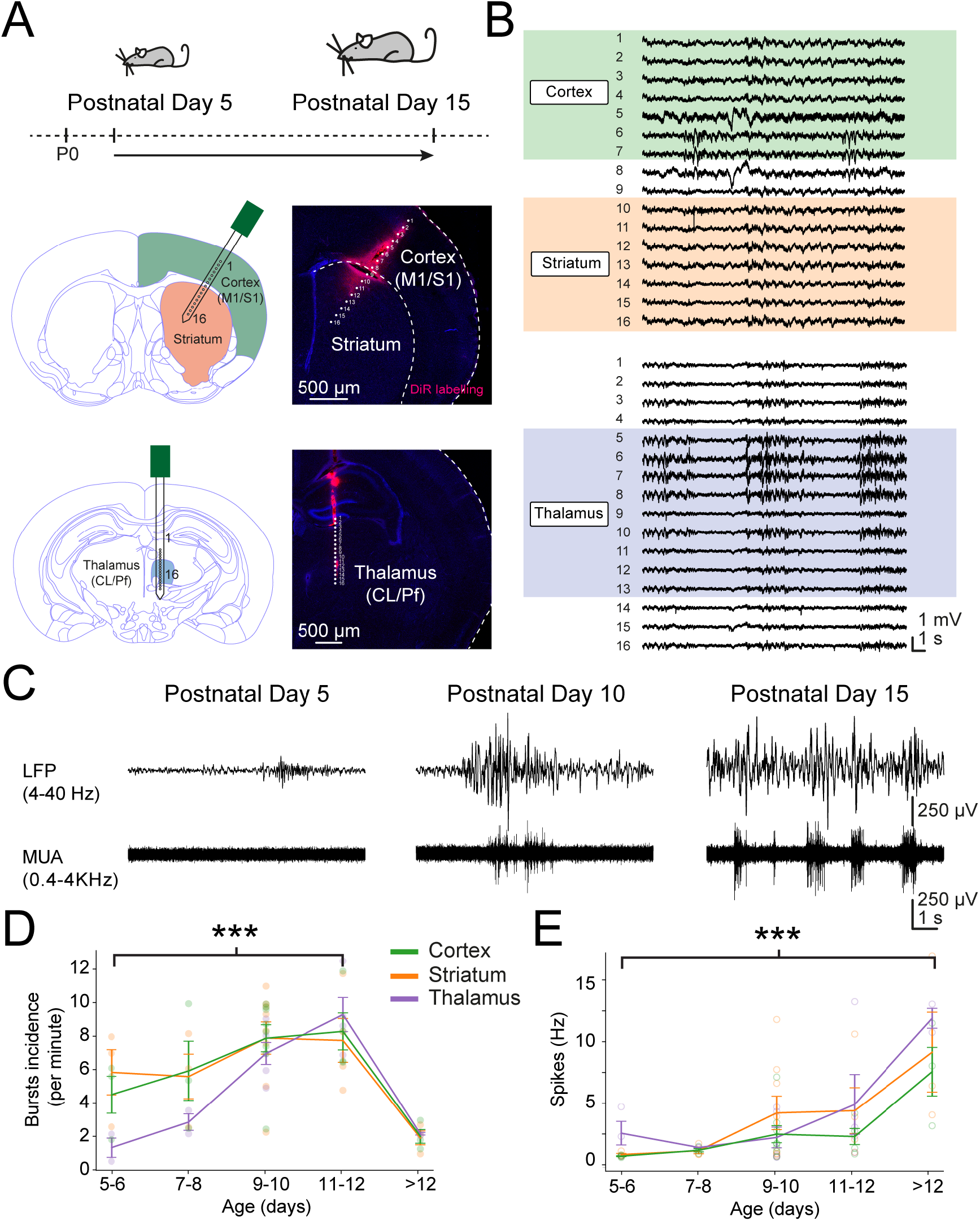
Overall neural activity in the basal ganglia circuit increases across early postnatal development. (**A**) Recordings were made from the cortex (M1/S1), dorsal striatum and intralaminar thalamus of mice between postnatal day (P)5 and P15 using a 16-channel silicon probe (5 mm - 100 µm channel spacing) inserted at a 30° angle to simultaneously record from cortex and striatum and an additional 16-channel probe (5 mm - 50 µm channel spacing) inserted vertically to record from the intralaminar thalamus, including the central lateral (CL) and parafascicular (Pf) nuclei. (**B**) Example field potential recordings made from the channels located in cortex and striatum (top) and thalamus (bottom) at P12. Note the heterogeneity in the amplitude, duration, and localisation of bursts. (**C**) Example striatal field potential recordings after 4-40Hz band-pass filtering to resolve bursts (top) and 0.4-4KHz band-pass filtering to resolve spiking or multi-unit activity (MUA, bottom). Note the general transition from small infrequent bursts interspersed by periods of silence to continuous activity patterns in the local field potential concomitant with an overall increase in spiking activity across early postnatal development. (**D**) Overall burst incidence modestly increased during early postnatal development. During later stages bursts were replaced with continuous oscillatory activity as observed in all three brain regions. (**E**) MUA increased in all three brain regions and was most pronounced during later stages of postnatal development.

### Unbiased classification and clustering of network bursts in cortical recordings

Initial inspection of bursting events suggested that they fell into roughly two distinct classes based on the dominant frequency of their oscillations and these could often be observed in recordings from all three brain regions (Figure 2). One class of events was reminiscent of spindle bursts (SB) (Figure 2A), due to their prominent power in the spindle frequency range (8-30 Hz) (Hanganu *et al*., 2006; Yang *et al*., 2016) (Figure 2B, C), while the second class of events was reminiscent of gamma bursts (GB) (Figure 2A), due to the appearance of higher frequency beta-low gamma oscillations (20-40Hz) (Figure 2B, C). We observed that the latter was often nested within lower frequency spindle-band activity, as has also been observed in prefrontal and prelimbic cortex and other regions (Brockmann *et al*., 2011; Khazipov *et al*., 2013; Yang *et al*., 2013; Cichon *et al*., 2014; Hartung *et al*., 2016a) (Figure 2B, C). In addition to these spectral characteristics, the GB often appeared longer in duration and exhibited increased MUA as compared to SB (Figure 2D, E). Due to the similarities between SB and GB events, e.g., both featuring elevated power at spindle frequencies and co-occurring at similar developmental time points, and to avoid risk of experimenter bias — for example, towards known patterns of bursting activity based on the existing literature, the burst events were categorized quantitatively using a combined principal component analysis (PCA) and clustering approach (see **Methods**). In brief, each detected bursting event was used to produce a feature vector incorporating several parameters including burst duration, power in the theta-alpha and beta-low gamma frequency range and spike rate, amongst others. By applying PCA, these features could be mapped to a lower dimensional space and then segmented via a clustering algorithm to identify different classes of bursting events. We first validated this approach on our cortical recordings to investigate whether it allowed for detection of similar events as previously described in the literature (Khazipov *et al*., 2004; Hanganu *et al*., 2006; Yang *et al*., 2009; Brockmann *et al*., 2011; Yang *et al*., 2013; An *et al*., 2014; Cichon *et al*., 2014; Shen & Colonnese, 2016). Using this approach, it was possible to capture 79% of the total variance across the feature space in the first three PCA components (Supplemental Figure 3A).

**Figure 2:**
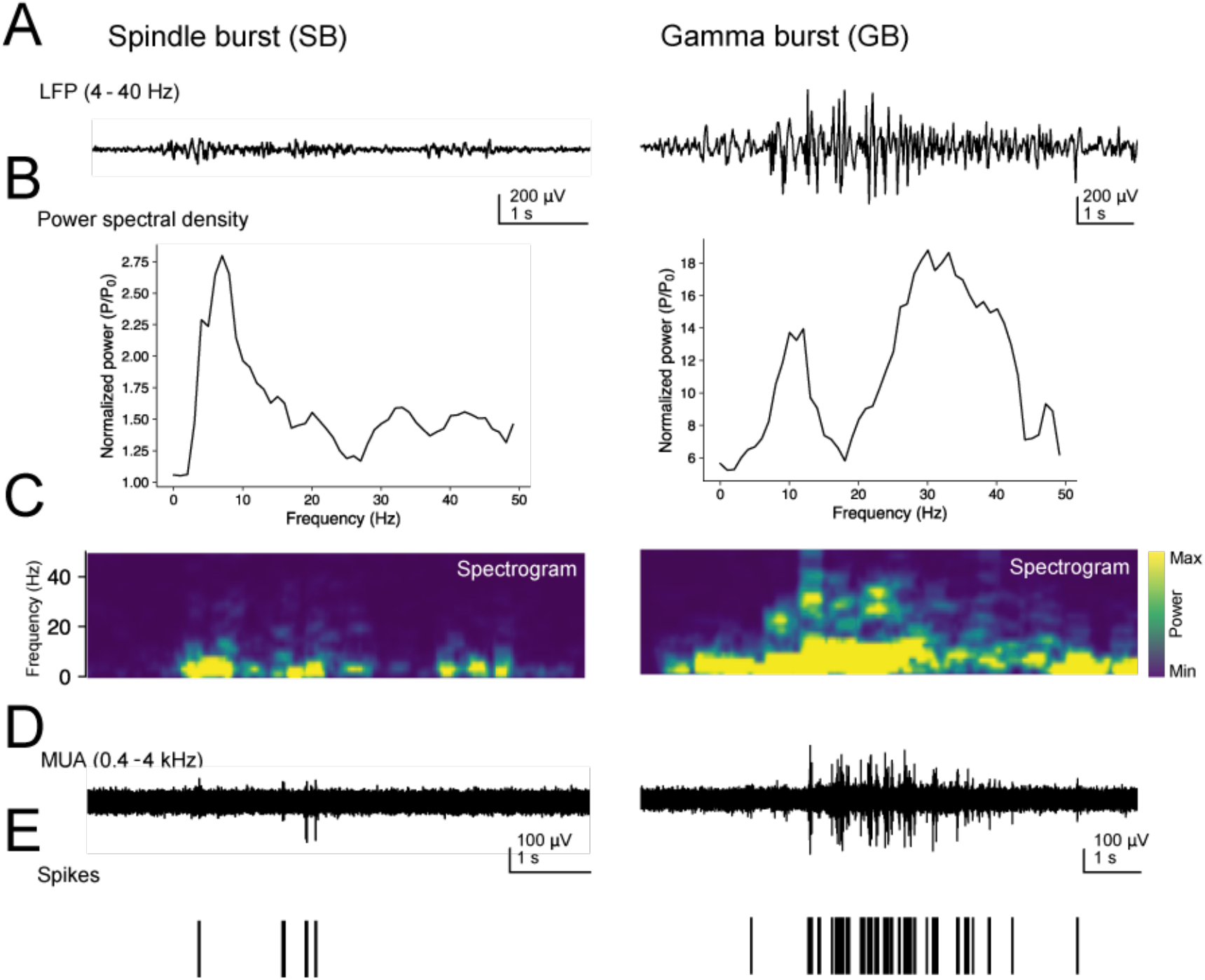
The main bursts events observed in the developing basal ganglia resemble spindle bursts and gamma bursts. (**A**) Example striatal spindle burst (SB) and gamma burst (GB) in 4-40 Hz bandpass filtered LFP signal (top) as recorded at postnatal day (P)8 and P10 respectively. (**B**) Power spectral density plot (PSD) of relative spectral power within bursts, normalized to baseline power. Note the prominent peaks at 5-10 Hz in both bursts and the additional peak at 20-40 Hz in the GB. (**C**) Spectrogram of the corresponding LFP signal. Note the presence of beta-gamma-band activity during the mid-phase of the GB. (**D**) 0.4-4 kHz MUA filtered activity. (**E**) Raster plot of spikes extracted from MUA. Note that GB are characterised by increased spiking at higher frequencies as compared with SB.

Subsequent clustering revealed two main classes of burst activity in our recordings from the motor/somatosensory cortex, which we refer to as SB and nested gamma spindle bursts (NGB) (Figure 3A). These exhibited characteristics similar to SB and GB previously described in several cortical regions (Hanganu *et al*., 2006; Yang *et al*., 2009; Cichon *et al*., 2014; Hartung *et al*., 2016a; Yang *et al*., 2016) and included similar dominant frequency components (Figure 3B, C) with a peak for SB at 4-16 Hz, referred to as theta-alpha (8-α) frequency, and for NGB an additional faster frequency peak at 20-30 Hz, referred to as beta-low gamma (β-ψ_Low_) frequency, as well as other corresponding parameters (Figure 3D and Supplemental Table 1 **and** 4). The NGB were very often nested within an envelope of SB frequency oscillations and are reminiscent of those described in the prefrontal cortex where they are referred to as NG (Brockmann *et al*., 2011). Importantly, we find that these detected cortical burst events exhibit a similar developmental pattern of changes as previously observed (Hanganu *et al*., 2006; Brockmann *et al*., 2011; An *et al*., 2014; Cichon *et al*., 2014; Shen & Colonnese, 2016). In particular, these include a progressive decrease in the incidence of SB events over developmental time while the incidence of NGB events increased (SB: 4.6 ± 2.1 to 2.4 ± 2.6 bursts/minute and NGB: 0.6 ± 0.6 to 4.9 ± 1.2 bursts/minute, F(3, 30) = 3.35, p = 0.0318, Figure 3E). We find that SB events were overall smaller in amplitude (SB: 502 ± 57.8 and NGB: 1040 ± 231 µV, F(1, 28) = 66.0, p = 7.59e-09) and were shorter in duration (SB: 1.34 ± 0.14 and NGB: 4.19 ± 1.09 s, F(1, 28) = 109, p = 3.73e-11) than NGB events. Both types of event exhibited a progressive developmental increase in their amplitude (SB: 356 ± 37 to 731 ± 48 µV and NGB: 993 ± 352 to 1282 ± 259 µV, F(1, 28) = 5.60, p = 0.00389) and duration (SB: 0.9 ± 0.1 to 1.5 ± 0.2 s and NGB: 3.1± 1.2 to 5.1 ± 1.2 s, F(1, 28) = 5.59, p = 0.00391, Figure 3E). Whereas the relative theta-alpha power remained constant throughout postnatal development for all burst events (SB: 0.3 ± 0.01 to 0.2 ± 0.02 A.U. and NGB: 0.3 ± 0.05 to 0.3 ± 0.02 A.U., Supplemental Figure 7Ai), the peak theta-alpha frequency slightly slowed for NGB events from ∼12 Hz at P5-6 (F(1, 28) = 5.35, p = 0.0282) to ∼9 Hz at P11-12 converging to the peak theta-alpha frequency of SB events (F(1, 28) = 8.30, p = 0.000419, Supplemental Figure 7Aii). As expected the relative beta-low gamma power was significantly greater for NGB events throughout development (SB: 0.3 ± 0.03 to 0.3 ± 0.03 A.U. and NGB: 0.5 ± 0.08 to 0.4 ± 0.04 A.U., F(1, 28) = 77.5, p = 1.48e-09, Supplemental Figure 7Aiii), although this did reduce slightly with age (F(3, 28) = 3.26, p = 0.0362), and the peak beta-low gamma frequency remained mostly constant at ∼22 Hz throughout development, although a trend towards faster frequencies was seen at later developmental stages (F(1, 28) = 2.93, p = 0.0509, Supplemental Figure 7Aiv).

**Figure 3:**
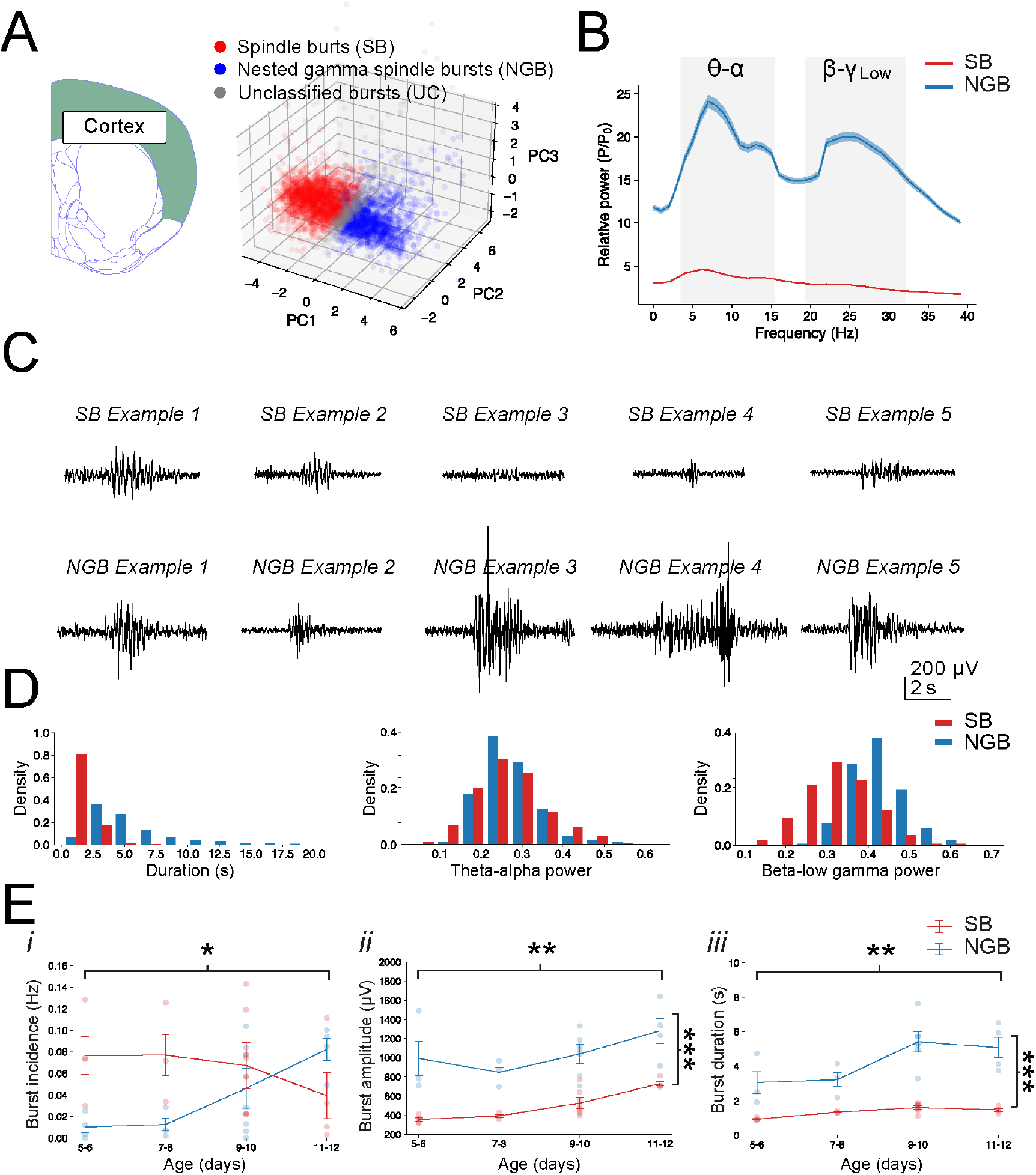
Unbiased detection and clustering of cortical bursts confirms presence of two main classes of bursts whose properties shift developmentally. **(A)** Scatter plot of the first three principal components used for clustering of cortical bursts into spindle bursts (SB, total number of 2237 events, red) and nested gamma spindle bursts (NGB, 1657 events, blue). Unclassified bursts (UC, 431 events) are denoted in grey. (**B**) Power spectral density (PSD) plot of the mean normalized power across SB and NGB events demonstrate that SB events have prominent power in the spindle frequency range (theta-alpha (8-α), 4-16 Hz), while NGB exhibit prominent power in the beta-low gamma frequency (β-ψ_Low_, 20-30 Hz) which are nested within lower frequency spindle-band activity. (**C**) Example of recorded cortical SB and NGB events after automated detection and clustering. Note the larger amplitude and longer duration of NGB events. (**D**) Histogram of the distributions of several key features across cortical SB and NGB events, including their duration, theta-alpha (8-α) power and beta-low gamma (β-ψ_Low_) power. Note the longer durations of NGB events and their increased beta-low gamma (Ω-ψ_Low_) power. (**E**) Graphs depicting the developmental changes in the incidence (*i*), amplitude (*ii*) and duration (*iii*) of SB and NGB events. Note the decrease in incidence of SB events and concomitant increase in NGB events across development, as well as an increase in both the amplitude and duration of events with the latter most pronounced for NGB events during later stages of development.

Overall, the cortical bursts detected and classified here exhibit all the hallmarks of those previously described in various cortical regions (Hanganu *et al*., 2006; Brockmann *et al*., 2011; An *et al*., 2014; Cichon *et al*., 2014; Shen & Colonnese, 2016) and suggests this approach of detection and unbiased classification could be effective in detection of burst events in other brain regions also.

### Detection of burst events in dorsal striatum and intralaminar thalamus

We next employed this detecting and clustering method on our striatal (Figure 4A) and thalamic recordings (Figure 5A). The first three components explained 80% and 76% of total variance across the feature space for respectively striatal and thalamic burst events (Supplemental Figure 3B, C).

**Figure 4:**
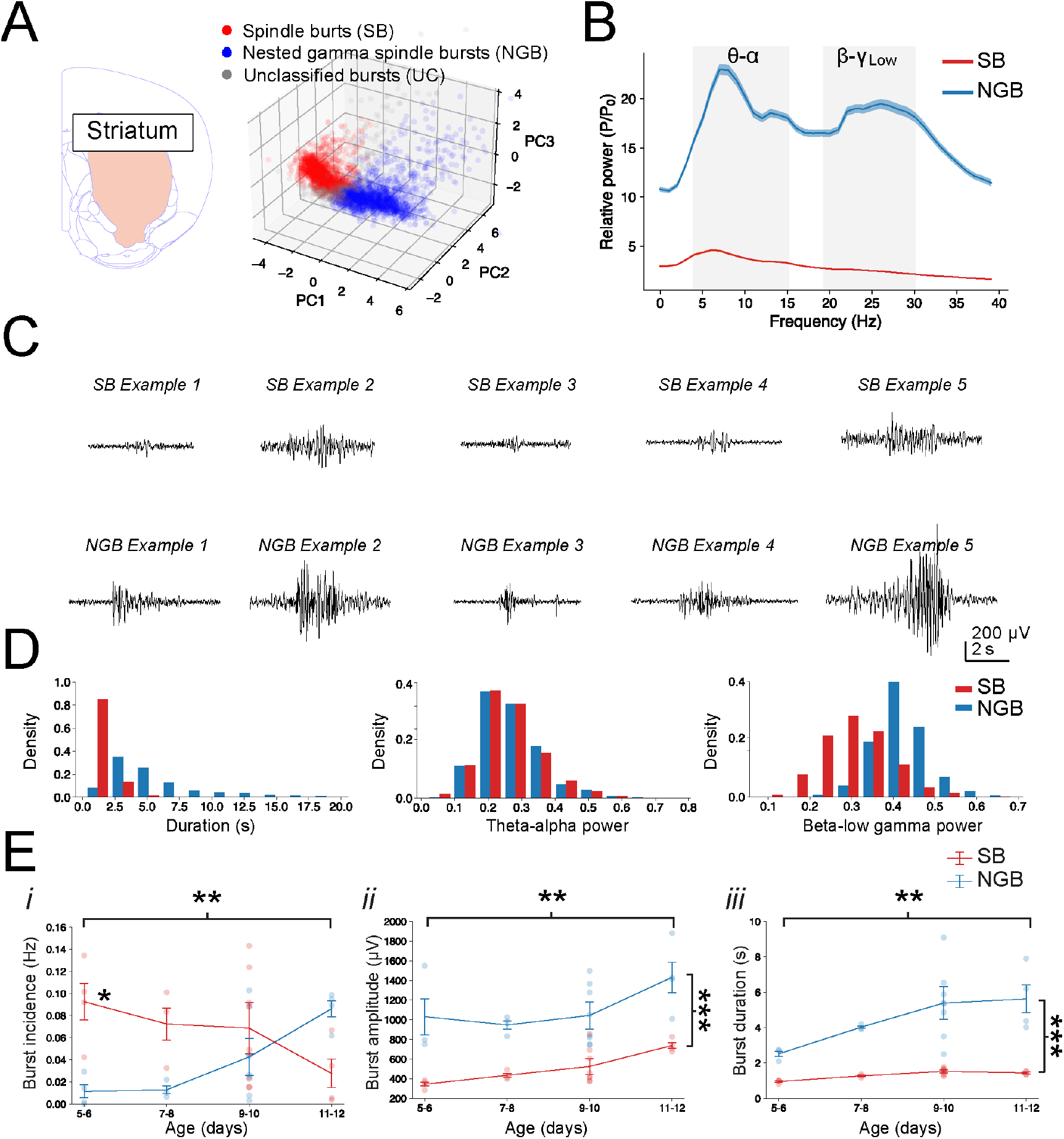
Striatal bursts also cluster in two groups whose properties are consistent with SB and NGB events. (**A**) Scatter plot of the first three principal components used for clustering of striatal bursts into spindle bursts (SB, total number of 2242 clustered events, red) and nested gamma spindle bursts (NGB, 1633 events, blue). Unclassified bursts (UC, 437 events) are denoted by grey (left). (**B**) PSD of the mean normalized power across striatal NGB and SB events reveals that both events exhibited a prominent peak at theta-alpha frequency (8-α) and that NGB events contain an additional peak at beta-low gamma frequency (β-ψ_Low_). (**C**) Example SB and NGB events in striatum. Note the larger amplitude of NGB events. (**D**) Histogram of the distributions of several key features across striatal SB and NGB events. (**E**) Graphs depicting the developmental changes in the striatal burst incidence (*i*), amplitude (*ii*) and duration (*iii*). Note the overall higher incidence of SB events (1,30) = 4.75, p = 0.0373) and their decrease and concomitant increase in NGB events across development, as well as an overall increase in the amplitude (F(3,29) = 4.73, p = 0.00831) and duration (F(3,29) = 3.93, p *=* 0.0181) of events with the latter most pronounced for NGB events.

**Figure 5:**
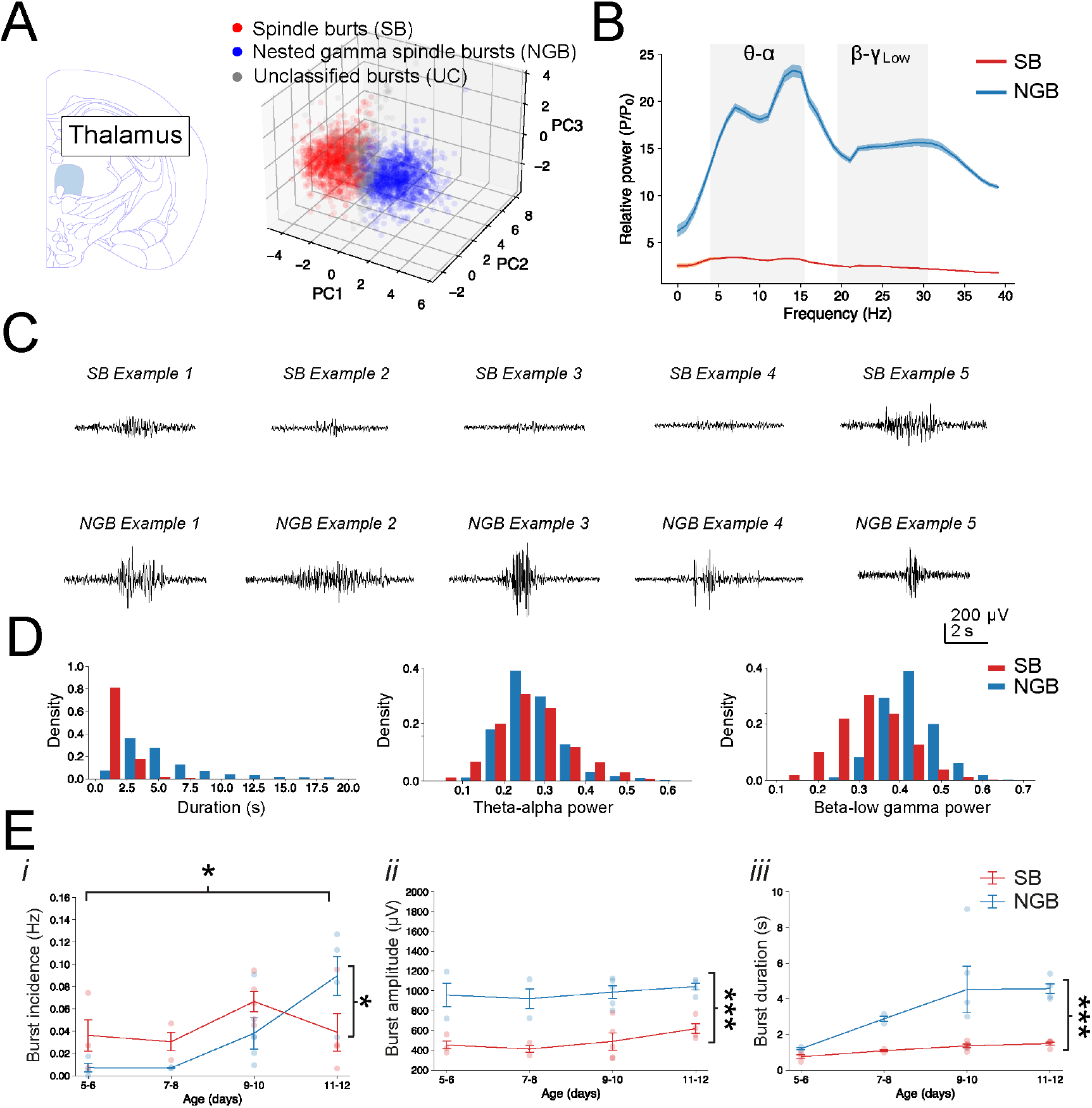
Thalamic bursts also cluster in two groups whose properties are consistent with SB and NGB events. (**A**) Scatter plot of the first three principal components used for clustering of thalamic bursts into SB (1384 events, red), NGB (1335 events, blue) and UC bursts (409 events, grey, left). (**B**) PSD of the mean normalized power across SB and NGB events. (**C**) Examples of SB and NGB events in thalamus. (**D**) Histogram of the distributions of several key features across thalamic NGB and SB events. (**E**) Developmental changes in the burst incidence (*i*), amplitude (*ii*) and duration (*iii*). Note the increase in thalamic NGB events across development and their consistent larger amplitude and longer duration.

Interestingly, the detected and clustered events in both striatum and thalamus exhibited many similar properties of those previously observed in cortex and it was opted to refer to these striatal/thalamic events also as SB and NGB events. This was guided by several observations. Firstly that, as in cortex, the relative oscillatory power of SB and NGB events differed in that SB events had an overall lower oscillatory power as compared to NGB. Secondly, that the mean power spectral density (PSD) across SB and NGB events in both striatum and thalamus all exhibited a peak at the spindle frequency centred at 5-10Hz (theta-alpha, 8-α) and NGB events exhibited an additional secondary peak centred at the 20-30Hz (beta-low gamma, β-ψ_Low_) range (Figure 4B, 5B **and** Supplemental Table 2, 3 **and** 5, 6). Beyond these spectral measures, NGB events in both striatum and thalamus were also significantly longer in duration and had a greater spike rate than SB events, again confirming our initial observations in cortex (Figure 4B, 5B **and** Supplemental Table 2, 3 **and** 5, 6). Moreover, their developmental properties were similar to those observed in cortex, including changes in event incidence with NGB incidence increasing with age at the cost of SB incidence (striatum: F(3,30) = 5.83, p = 0.00290 and thalamus: F(3,20) = 3.50, p = 0.0345, Figure 4E and 5E), as well as a consistently larger amplitude of NGB events (striatum: (1,29) = 55.0, p = 3.59e-08 and thalamus: (1,19) = 57.77, p = 3.56e-07, Figure 4E and 5E) and duration (striatum: F F(3,29) = 3.93, p *=* 0.0181 and thalamus: (1,19) = 20.96, p = 0.000205, Figure 4E and 5E). In addition, many other changes in spectral properties of events mirrored that seen in cortex (Supplemental Figure 7). Finally, a number of events in all three brain regions whose properties did not fit either category sufficiently, and often had characteristics in between those of SB and NGB, were deemed unclassified (Supplemental Figure 4). Taken together these results suggest that the early network activity in the striatum and thalamus are also dominated by two distinct classes of network bursts, namely spindle bursts and nested gamma spindle bursts.

### Region-specific properties of network bursts

To further assess how the SB and NGB events differed across brain regions a multivariate extension of the F-ratio test was performed to determine whether the distributions of features differed across each class of burst events between the different brain regions (see **Methods**). This analysis revealed that many of the properties of SB and NGB events exhibited region-specific differences (Supplemental Figure 5). For example, although SB events were similar between striatum and thalamus (F(1, 3624) = 6.82, adjusted p = 0.0711), their distribution of features significantly differed between cortex and thalamus (F(1, 3619) = 30.2, adjusted p < 0.001) as well as between cortex and striatum (F(1, 4477) = 53.8, adjusted p < 0.001; Supplemental Figure 5A, C, E). For NGB events it seemed that cortex and striatum were more similar (F(1, 4477) = 2.13, adjusted p = 1.0), whereas NGB events differed between cortex and thalamus (F(1, 2990) = 30.8, adjusted p < 0.001) and between striatum and thalamus (F(1,2966) = 22.8, adjusted p < 0.001; Supplemental Figure 5B, D, F).

These findings indicate that the variation in the feature distributions between SB and NGB events in some, but not all regions, exceeded the levels expected by chance. For comparison, the equivalent F-ratios between SB and NGB events within each brain area were several orders larger (cortex: F(1, 3892) = 8461, adjusted p < 0.001; striatum: F(1, 3873) = 8510, adjusted p < 0.001; thalamus: F(1, 2717) = 5464, adjusted p < 0.001; Supplemental Figure 5G, H, I). Thus, even where variation exceeded significance between certain brain regions for the same class of events (e.g. striatal versus thalamic NGB events), these differences were considerably smaller compared with variation between different classes of burst events within one brain region.

As a *post hoc* analysis to dissect how events differed across brain regions, we performed pairwise comparisons along each feature dimension (e.g. duration of burst events). Due to the extensive sample size, most comparisons reached significance even after correcting for multiple comparisons (Supplemental Table 7 **-** 12). However, the effect sizes between events were mostly small (range = 0.10-1.05) indicating that differences were driven by differences across spectral and structural features rather than any individual dimension. Nonetheless some key differences were observed. For example, that for striatal and cortical NGB events the peak in the theta-alpha (8-α) range, similar to cortex, was centred around ∼5 Hz with significantly greater power in the 4-12 Hz range over the 12-20 Hz range (Welch’s t(2983) = 4.91, p = 9.60e-07), whereas for thalamic events the peak frequency was centred around ∼12 Hz with significantly greater power in the 12-20 Hz range over the 4-12 Hz range (Welch’s t(2517) = 6.32, p = 3.03e-10; Figure 3B, 4B and 5B). In addition, the highest spike rate was found in cortical and striatal NGB whereas in thalamus the highest spike rate was found for SB events (Supplemental Figure 6 **and** Supplemental Table 4, 5 **and** 6). Taken together these results suggest that the SB and NGB events exhibit subtle but significant differences between different brain regions which likely reflect the cellular environment of their recording and generation.

### Dynamic developmental changes in functional interactions between brain regions during bursting activity

We next determined how these bursts interacted amongst the three brain regions (Figure 6A). Firstly, it was investigated which brain regions had the most frequent co-occurring events by taking the proportion of events whose onset overlapped out of the total number of burst events across the paired regions.

**Figure 6:**
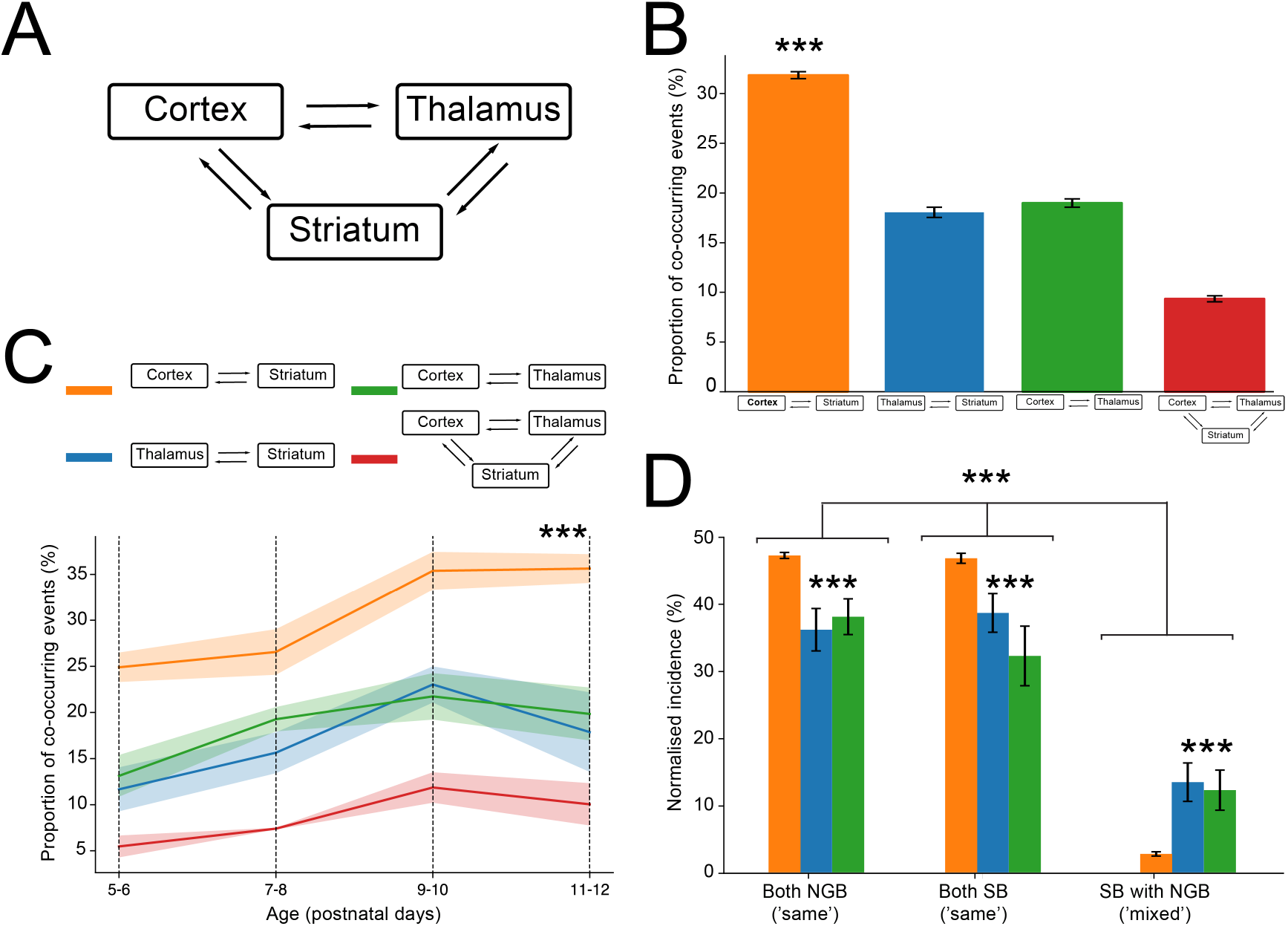
Co-occurring burst events are often of the same type and are increasingly observed during development. (**A**) The presence of co-occurring events was determined in the cortical, striatal, and thalamic regions based on their onset. (**B**) Co-occurring burst events are significantly more likely between cortex and striatum than between other brain regions. Only a small number of events co-occurred amongst all three brain regions (M = 0.09, SD = 0.04, all adjusted p ≤ 0.0134). (**C**) The proportion of co-occurring burst events significantly increases with postnatal age in all brain regions. Note that this is most pronounced for cortex-striatum between postnatal days 7-8 and 9-10. (**D**) Normalized incidence of NGB-NGB, SB-SB (both ‘same’) and NGB-SB (‘mixed’) events amongst different brain regions. ‘Same’ events are more likely than ‘mixed’ events, implying that communication across brain areas may be mediated by the same rather than different burst types.

This revealed that although all brain regions exhibited events that co-occurred, the incidence of co-occurring events was greatest (∼30%) between cortex and striatum (all adjusted p<0.001, Figure 6B). We next determined how these co-occurring bursting events changed across developmental time and find that all combinations of brain regions showed an increase in co-occurring events across developmental time (F(3, 41) = 8.50, p = 0.000164, Figure 6C). Lastly, we explored whether co-occurring bursting events are more likely to be of the ‘same’ class or different class (‘mixed’) of burst event (Figure 6D). There was a significant effect of burst type on the normalized incidence (F(2, 163) = 389, p = 9.59e-63), where the normalized incidence of co-occurring NGB/NGB events (M = 0.42, SD = 0.08) was significantly greater than co-occurring ‘mixed’ NGB/SB or SB/NGB events (M = 0.07, SD = 0.08; t(109) = 22.9, p = 1.36e-43, d = 4.36), as was the normalized incidence of co-occurring SB/SB events (M = 0.42, SD = 0.11) events over mixed events (t(109) = 19.4, p *=* 4.88e-37, d = 3.62). This appeared to be the case for all three brain region pairs – although the interaction effect of burst type and brain region also reached significance (F(4, 163) = 17.3, p = 7.29e-12), where co-occurring events appeared to differ in their type of event more often amongst cortex-thalamus and thalamus-striatum (Figure 6D). Lastly, there was no significant difference in the normalized incidence of NGB/NGB versus SB/SB events (t(110) = 0.54, p = 0.593, d = 0.10).

Together, these results indicate that co-occurring burst events are significantly more likely to belong to the same rather than different burst class, implying that communication across brain areas may be mediated by the same rather than different burst types.

Having identified that bursts of the same type are more likely to co-occur in different brain regions, next the cross-spectral coherence across pairs of NGB and SB events were analysed to quantify the extent to which events were similar between brain regions. Firstly, it was found that the mean cross-spectral coherence was significantly greater for both NGB and SB pairs compared to randomly shuffled data (p < 0.001, Supplemental Figure 8 and see **Methods**), indicating that the observed coherence is above that expected by chance. We then went on to explore the mean cross-spectral coherence across pairs of SB and pairs of NGB events. This revealed that the mean cross-spectral coherence in cortex-striatum was greatest for both SB (M = 0.61, SB = 0.11) and NGB events (M = 0.64, SD = 0.09) as compared to other brain region pairs (NGB and SB, all adjusted p ≤ 2.02e-24; Figure 7A). Within cortex-striatum the coherence among co-occurring events was greatest for NGB events (t(1219) = 5.87, p = 5.54e-09) (Figure 7A). In contrast, for thalamus-striatum events, coherence was significantly greater for SB (M = 0.48, SD = 0.11) over NGB events (M = 0.24, SD = 0.12; t(222) = −20.7, p = 7.95e-54), as was the case for cortex-thalamus (SB events: M = 0.45, SD = 0.13 and NGB events: M = 0.24, SD = 0.11; t(218) = −17.7, p = 5.12e-44, Figure 7A). This suggests that different brain regions preferentially synchronize during different types of bursts. Interestingly, burst events across different regions exhibited distinct changes in mean cross-spectral coherence over the course of development (Figure 7B). For events in cortex-striatum they appeared mostly stable in power over time and in fact slightly increased in coherence (NGB: 0.60 ± 0.12 to 0.65 ± 0.08; SB 0.59 ± 0.11 to 0.70 ± 0.07; F(3, 1610) = 15.8, p = 3.67e-10). In contrast, for events in thalamus-striatum coherence declined for both SB and NGB (NGB: 0.58 ± 0.10 to 0.23 ± 0.09; SB 0.54 ± 0.08 to 0.49 ± 0.10; F(3, 520) = 27.1, p = 2.80e-16) with the largest decrease seen for NGB events (F(3, 520) = 8.78, p = 1.09e-05). A similar trajectory could be observed for events in cortex-thalamus with a similar an overall decline in cross-spectral coherence during development (NGB: 0.52 ± 0.12 to 0.24 ± 0.10; SB 0.53 ± 0.07 to 0.46 ± 0.10; F(3, 538) = 23.5, p = 2.71e-14) which was markedly greater for NGB events (F(3, 538) = 4.58, p = 0.00354).

**Figure 7:**
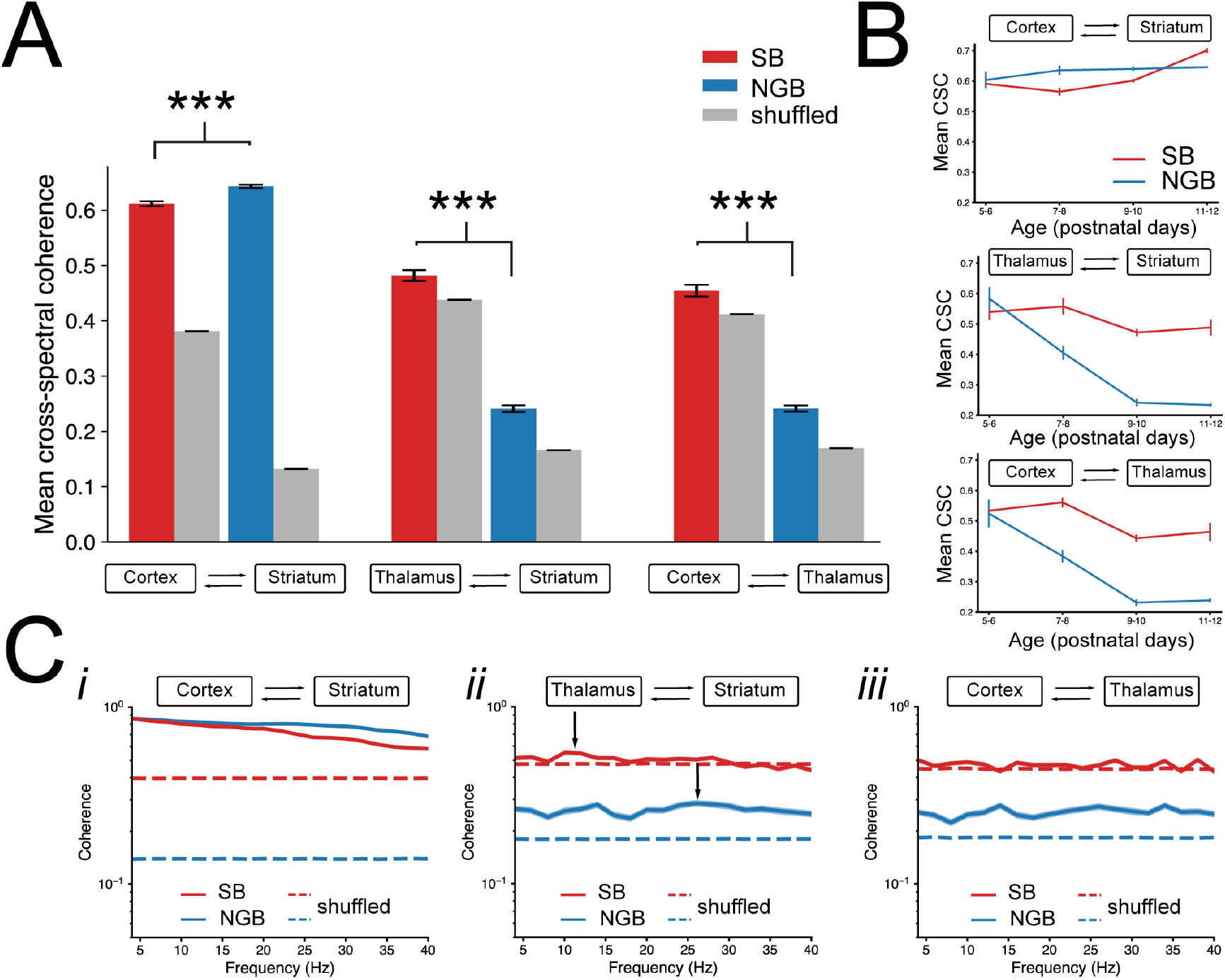
The coherence is greatest between cortex and striatum during NGB events. (**A**) The mean cross-spectral coherence is larger for NGB than SB for corticostriatal interactions. In contrast the mean cross-spectral coherence is larger for SB bursts event in both thalamostriatal and corticothalamic interactions. (**B**) Mean cross-spectral coherence across developmental time remained constant for corticostriatal events but significantly declined for NGB events in thalamostriatal and corticothalamic interactions. (**C**) Coherence analysis in the 4-40Hz frequency bands for corticostriatal events (*i*), thalamostriatal events (*ii*) and corticothalamic events (*iii*). Note the broad coherence across frequencies and significance for both SB and NGB events for cortex-striatum but only for NGB events in other brain regions.

We next investigated whether certain frequencies within co-occurring bursts of the same type exhibited greater coherence than others and analysed the coherence spectra within the 4-40 Hz frequency range to capture the dominant frequencies of both SB and NGB events (Figure 7C). This revealed that for cortex-striatum events, the cross-spectral coherence was significantly greater than that for shuffled data across the full range of frequencies 4-40 Hz for both SB and NGB events (Figure 7Ci). In contrast, for thalamus-striatum interactions, while NGB events were significant across all frequency ranges (4-40 Hz), there was a reduction in coherence for SB events as a function of frequency which did not maintain significance above ∼30 Hz, indicating that SB events exhibit some coherence at lower frequencies (Figure 7Cii). Lastly, for cortex-thalamus events, co-occurring NGB bursts were above the significance threshold across all frequencies, while SB events showed a more mixed pattern dipping below threshold around ∼14Hz as well as ∼36 Hz (Figure 7Ciii). These observations were consistent with those obtained using imaginary coherence analyses, which suppresses zero-lag coherence, which suggested that NGB-NGB co-occurring events are truly coherent across regions, but also that it is not possible to exclude the possibility that SB-SB co-occurring events are impacted by volume conductance (Supplemental Figure 9).

To determine whether one brain region was driving the other during these synchronous bursting events, we next performed a cross-correlation analysis to calculate the possible lags between co-occurring burst events (see **Methods** and Figure 8A, B). Frequency bands of interest were taken as both 4-16 Hz and 16-40 Hz ranges, around which spectral power was greatest for respectively SB and NGB bursts. To maximize statistical power due to lower numbers of co-occurring events, we split the data into two groups roughly corresponding to the first postnatal week (P5-8) and second postnatal week (P9-12). Overall, we find that the interaction between cortex and striatum produced the largest cross-correlation values irrespective of age or type of burst, indicating that activity between these brain regions is highly correlated (P5-8 SB max ρ^2^ = 0.00675, P5-8 NGB max ρ^2^ = 0.0297, P9-12 SB max ρ^2^ = 0.0256, P9-12 NGB max ρ^2^ = 0.0243, Figure 8). However, cortex-thalamus also exhibited large cross-correlations at P5-8 in the 16-40 Hz range (P5-8 SB max ρ^2^ = 0.00525, P5-8 NGB max ρ^2^ = 0.0189, P9-12 SB max ρ^2^ = 0.00175, P9-12 NGB max ρ^2^ = 0.00148, Figure 8).

**Figure 8:**
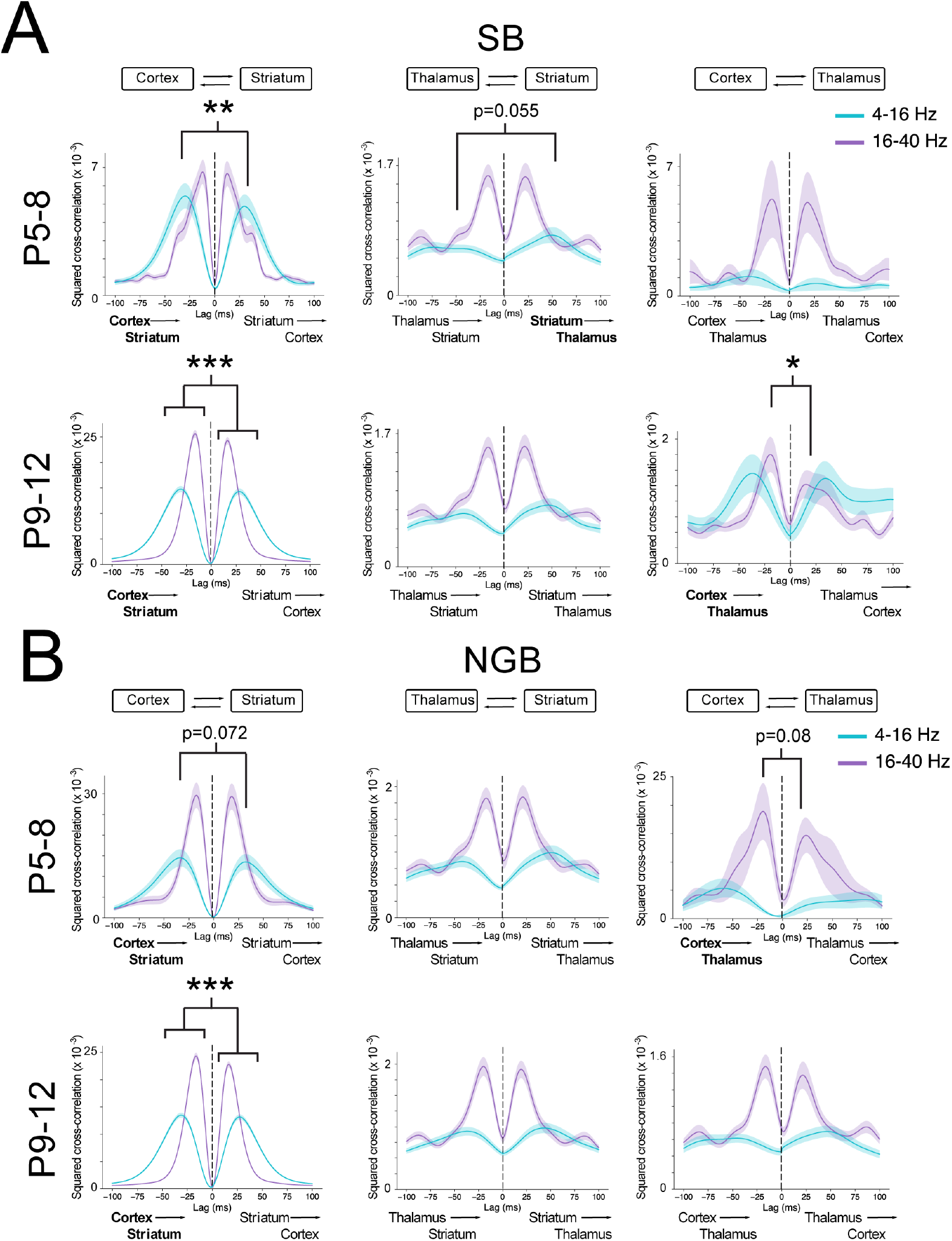
Cortical activity mainly drives activity in downstream basal ganglia. (**A**) Results from cross-correlation analysis for SB events that occur between P5-8 (top) and between P9-12 (bottom). Note the significant lead of cortex driving striatal SB events at P5-8 specifically at lower frequencies (4-16 Hz). At later developmental stages cortex significantly leads striatum in both frequency bands. (**B**) Results from cross-correlation analysis for NGB events. Note the trend for cortex to lead both striatum and thalamus at early developmental stages (P5-8, top) but cortex leads striatum significantly only at later developmental stages (P9-12, bottom).

Between the various brain region pairs, there was a clear peak in the 16-40 Hz band centred around +19/-17 ms, consistent with putative monosynaptic interactions amongst all regions (Hartung *et al*., 2016) (Figure 8). In contrast, the 4-16 Hz band was characterised by a broader cross-correlation curve centred around +41/-42 ms. Using this analysis to explore for potential lags between co-occurring burst events we find evidence of a significant cortical lead across both cortex-striatum and cortex-thalamus interactions, implying a potential cortical origin to these subcortical bursting events. For cortex-striatum events there was a significantly larger peak specifically in the 4-16 Hz range for SB events at −29 ms lag (Wilcoxon rank sum test, p = 0.00293), indicating a putative cortical lead for SB events during early stages of development (P5-8) (Figure 8A) which was not seen for NGB events at either frequency range. At later developmental stages (P9-12) we find a more universal significant negative lag across both frequency bands for both SB and NGB events (SB: 4-16 Hz, −16 ms; SB: 16-40 Hz, −31 ms; NGB: 4-16 Hz, −16 ms; NGB: 16-40 Hz, −31 ms; Wilcoxon rank sum test, all p ≤ 0.000128), indicating a more general cortical lead of activity in the striatum (Figure 8A, B). For cortex-thalamus events there was a trend towards a cortical lead at P5-8 for NGB events in the 16-40 Hz band with a −20 ms lag, though this comparison did not reach significance (Wilcoxon rank sum test, p = 0.0803) but at later developmental stages we find evidence of a cortical lead at P9-12 for SB events at −19 ms lag (Wilcoxon rank sum test, p = 0.0406, Figure 8A). Finally, there was no evidence for a significant lead in the peaks for thalamus-striatum events (Wilcoxon rank sum test, all p > 0.05), suggesting that neither region clearly drives burst events (Figure 8A, B). Together, these results suggest that at different stages of early postnatal development different parts of the basal ganglia network communicate with each other preferentially at different frequencies with the strongest interaction between cortex and striatum over and above interactions amongst other brain regions. The observations were also consistent with those obtained using spike-field coherence analysis (Supplemental Figure 10) and spike-density cross-correlation analysis (Supplemental Figure 11), which are less sensitive to volume conduction issues.

## Discussion

Here we describe for the first time the developmental bursts of activity that occur within key nodes of the neonatal mouse basal ganglia, including the cortex and intralaminar thalamus input structures and the dorsal striatum. Using an unbiased detection and classification approach we find that network bursts can be broadly classified into two different types and include spindle bursts (SB) and nested gamma spindle bursts (NGB), which are observed in all three brain regions. Analysis of these bursts across postnatal days (P)5 to P12 suggests that these patterns of activity exhibit several developmental changes including a developmental decrease in SB events with a concomitant increase in NGB events, which also become progressively longer in duration. Many of the bursts occurred synchronously in the different basal ganglia regions and were often, but not always, of the same type. This co-occurrence of events exhibits a developmental increase and was most pronounced between events in cortex and striatum. Indeed, the coherence is also greatest between cortex and striatum and in particular during NGB events. In contrast, the coherence between thalamus and striatum and cortex and thalamus is greatest during SB events. Cross-correlation analysis suggests that the high coherence reflects cortical activity driving striatum during both the first and second postnatal weeks, initially preferentially at lower frequencies (4-16 Hz) and during the second postnatal week at higher frequencies (16-40 Hz). In addition, cortex also appears to drive activity in the thalamus and preferentially at higher frequencies (16-40 Hz) and mainly during later stages of development. Taken together these results demonstrate the dynamic nature of burst generation within the developing basal ganglia and aims to facilitate future explorations of the physiological functions of these bursts for its development.

We recorded network bursts in urethane anaesthetised C57 mice between P5-12 using multi-channel silicon probes. Recordings were made under urethane anaesthesia which is thought to help preserve naturalistic activity dynamics (Sitdikova *et al*., 2014; Shumkova *et al*., 2021) akin to a sleep state and so reflective of the default state of pups during this developmental period (Bolles & Woods, 1964; Clement *et al*., 2008). In general neonatal bursts are typically classified based on their structural and spectral properties and in developing cortex include ‘spindle bursts’ as recorded in rodents (Khazipov *et al*., 2004; Hanganu *et al*., 2006; Yang *et al*., 2016), and their human equivalent ‘delta-brushes’ as recorded from preterm human neonates (Vanhatalo & Kaila, 2006; Milh *et al*., 2007; Vecchierini *et al*., 2007). These are thought to be one of the earliest activity patterns generated in the postnatal brain - see for early prenatal activity patterns in thalamus (Moreno-Juan *et al*., 2017; Anton-Bolanos *et al*., 2019), and consist of short bursts of activity containing 4-16 Hz oscillations which are often initiated by distal muscle twitches and are thought to contribute to the establishment of circuits required for sensorimotor coordination (Khazipov *et al*., 2004; Milh *et al*., 2007; Dooley & Blumberg, 2018). In addition, a second major pattern of activity, so-called ‘gamma bursts’ also appears transiently in cortex during the first postnatal weeks (Minlebaev *et al*., 2011; Khazipov *et al*., 2013; Yang *et al*., 2013) and consist of short bursts of activity in the gamma frequency range (20-40Hz). Both patterns of activity can result from peripheral activity such as whisker activation, muscle twitches or retinal waves and are often topographically organized according to the site of sensory stimulation (Yang *et al*., 2013; Gerasimova *et al*., 2014), but can also arise spontaneously from the intrinsic dynamics within the developing circuits (Hanganu *et al*., 2006; Yang *et al*., 2013; Luhmann *et al*., 2016; Martini *et al*., 2021). The early transient bursts of activity largely disappear at later stages of development and are replaced by persistent oscillatory activity, such as gamma oscillations (Khazipov *et al*., 2013). Finally, these early bursting patterns often do not occur in isolation but rather form complex, nested motifs. For example, transient gamma oscillations often precede spindle bursts (Yang *et al*., 2013) or can be nested within spindle bursts (Brockmann *et al*., 2011), and both spindle bursts and gamma oscillations might be nested within larger delta waves (Khazipov *et al*., 2013).

Overall, the literature on neonatal bursting activity is highly diverse, even when restricted to studies in rodent models and similar brain regions (Yang *et al*., 2009; Seelke & Blumberg, 2010; Brockmann *et al*., 2011; An *et al*., 2014). This diversity of findings likely results from a combination of factors, including exact recording protocols, differing levels of anaesthetics and/or recording ages, both species-specific and exact recording location-specific factors, and the lack of standardisation for classifying and naming of bursting events. We therefore opted for an unbiased detection and classification approach for our recordings. Using this approach, we suggest that two distinct classes of bursting activity can be reliably identified across cortex, striatum and thalamus and include spindle bursts (SB) and nested gamma spindle bursts (NGB) which exhibited properties very similar to those described previously in cortical regions, including their spectral/oscillatory properties with a peak for SB at 4-16 Hz and for NGB an additional faster frequency peak at 20-30 Hz (Contreras *et al*., 1997; Khazipov *et al*., 2004; Hanganu *et al*., 2006; Yang *et al*., 2009; Yang *et al*., 2013; An *et al*., 2014; Cichon *et al*., 2014; Gerasimova *et al*., 2014; Hartung *et al*., 2016a; Bitzenhofer *et al*., 2017; Murata & Colonnese, 2018). The bursts detected in our recordings were in general very similar to those described previously in cortical structures. For example, motifs of bursting activity in the prefrontal cortex of rodents are seen with similar discontinuous oscillatory events centred on spindle and beta-gamma frequencies (Cichon *et al*., 2014; Chini *et al*., 2020). However, subtle differences were observed in the properties of events in our recordings from motor/somatosensory cortex and those described in other cortical structures. For example, the duration of NGB events in our cortical recordings were predominantly longer than SB events, similar to those seen in more frontal cortical regions (Brockmann *et al*., 2011), but differing from those recorded in sensory cortex where bursts of gamma oscillations were significantly shorter than SB events (Yang *et al*., 2009). Although it is largely unclear whether similar bursting events share a common underlying cellular mechanism of generation in different brain regions (Hanganu-Opatz *et al*., 2021) it is likely that differences in cortical cellular and circuit architecture across visual, somatosensory and prefrontal cortex (Allene *et al*., 2008; Katzel *et al*., 2011; Tasic *et al*., 2018) can contribute to these observed differences. The bursts recorded and analysed in this dataset were very similar between brain regions while also exhibiting subtle region-specific differences. For example, during NGB bursts the overall power of the beta-gamma oscillations was greatest in cortex, the spike rate was greatest in thalamus and peak frequency of theta-alpha oscillations differed between brain regions ranging from 7 Hz for cortex and striatum to 14 Hz in thalamus. Spiking during neonatal thalamic bursts consisted of ∼5-20 action potentials at a frequency of around 5-30 Hz which is different to that seen during adulthood (Lacey *et al*., 2007) and they likely reflect unique developmental patterns of activity as a result of neuronal and circuit maturational state. These differences also likely reflect region-specific network dynamics, overall levels of neural activity and precise positioning of current sinks and sources (Tanaka & Nakamura, 2019).

Analysis of the changes of these bursts across development suggests a few interesting developmental trajectories for SB and NGB events which were evident in all brain regions studied. Overall, the NGB events tended to increase in incidence while SB events reduced in incidence across development. These developmental trajectories are consistent with the wider literature where spindle activity tends to decline with age (Nakazawa *et al*., 2020), while gamma bursts (Seelke & Blumberg, 2010) and gamma-band activity more generally increases over time (Minlebaev *et al*., 2011; Bitzenhofer *et al*., 2020). These changes in events likely reflect the developmental changes in the neuronal circuit maturation to be able to sustain these faster frequency events for longer periods of time. Within the cortex, gamma-band activity depends on interactions between interneurons – in particular perisomatic-targeting parvalbumin-positive cells — and pyramidal cells, where the reciprocal excitatory and inhibitory interactions are thought to give rise to gamma frequency oscillations (Tiesinga & Sejnowski, 2009). Thus, the developmental increase in gamma-band activity likely reflects the integration of perisomatic interneuron-mediated inhibition within glutamatergic circuits where earlier gamma bursts (pre-P5) are likely independent of GABAergic inhibition (Khazipov *et al*., 2013). However, what changes in cellular and circuit properties underlie the observed changes in oscillatory behaviour within the basal ganglia are largely unknown. Several different approaches could be employed to investigate when basal ganglia interneurons come ‘online’ by recording larger numbers of units and using waveform analysis to classify neuronal populations (Reyes-Puerta *et al*., 2015; Bitzenhofer *et al*., 2020). It might also be interesting to explore how the neonatal NGB bursts relate to the beta bursts seen in adulthood in cortical and basal ganglia circuits as they contain many of the same frequency components and might be subserved by similar circuits of interconnected neurons (Cagnan *et al*., 2019).

Analysis of the interactions amongst brain regions during burst activity revealed several interesting dynamical changes. Firstly, that events that co-occurred amongst brain regions often, but not always, were of the same type and that such interactions increased over developmental time. This likely reflects maturation of synaptic connections (Tepper *et al*., 1998; Nakamura *et al*., 2005; Dehorter *et al*., 2011; Peixoto *et al*., 2016; Krajeski *et al*., 2019). Secondly, that the coherence between brain regions was greatest between cortex and striatum and that this coherence was retained throughout the first postnatal week. This is consistent with the observation that in adulthood also the activity within striatum is strongly reflective of activity within cortex (Peters *et al*., 2021) as propagated by the various excitatory cortico-striatal afferents (Hunnicutt *et al*., 2016). The coherence between thalamus and striatum and cortex and thalamus was initially high also but this significantly lowered across development, suggesting progressive uncoupling or more local differential generation of these burst events. Thirdly, cross-correlation analysis of co-occurring bursts in different brain region would suggest that cortex leads activity in striatum initially at lower frequencies and later at faster frequencies. In addition, the cortex seems to lead activity in thalamus preferentially at faster frequencies. LFP signals arise from the summation of many aligned voltage sources and are expected to emerge more clearly from organized and laminar structures (e.g. cortex) than homogeneous structures such as the basal ganglia (Buzsaki *et al*., 2012). It is therefore possible to argue that such suggested interactions result from volume transmission. However, our analysis using imaginary coherence, spike-field coherence and spike-density cross-correlation would suggest that that some do reflect functional interactions as opposed to resulting from volume conduction only (Boraud *et al*., 2005) and in particular for NGB-NGB events in cortex and striatum. We did not observe a clear lead of activity between striatum and thalamus which suggests that activity might be led by a tertiary structure - for example, some common cortical area or areas. Indeed, as spindle bursts have been shown to propagate across diverse cortical regions (An *et al*., 2014) and even across hemispheres (Yang *et al*., 2009), the possibility of an extended cortical driver via long-range connections is plausible. Selective inactivation of certain brain regions (e.g. via lidocaine injection (Yang *et al*., 2013) or electrical or optogenetic stimulation (Bitzenhofer *et al*., 2017) would provide more conclusive evidence for which regions are driving others.

In conclusion, we find that the developing basal ganglia exhibits distinct oscillatory activity patterns during the first postnatal weeks, which can be classified as spindle bursts and nested gamma spindle bursts. These appear to be predominantly initiated in cortical regions and drive activity in downstream nuclei. While neural activity in general is necessary to drive maturation and survival of striatal neurons (De Marco Garcia *et al*., 2011; Kozorovitskiy *et al*., 2012; Peixoto *et al*., 2016; Peixoto *et al*., 2019; Sreenivasan *et al*., 2022) it is tempting to speculate that these different types of bursts have different functional roles during early development. Indeed, they might convey distinct developmentally relevant information to the basal ganglia as well as contribute towards the establishment of its fine-scale synaptic connectivity by engaging distinct synaptic plasticity rules (Kozorovitskiy *et al*., 2015; Valtcheva *et al*., 2017; Han *et al*., 2020; Mendes *et al*., 2020).

## Acknowledgements

This work is supported through an MRC Career Development Award (MR/M009599/1) and a John Fell Fund award (162/059) to TE. Oxford NDCN 4-year DPhil studentship to SKW. We thank Rohan Krajeski for initial help with the project.

## Supplemental Material

### Supplemental Figures

**Supplemental Figure 1:**
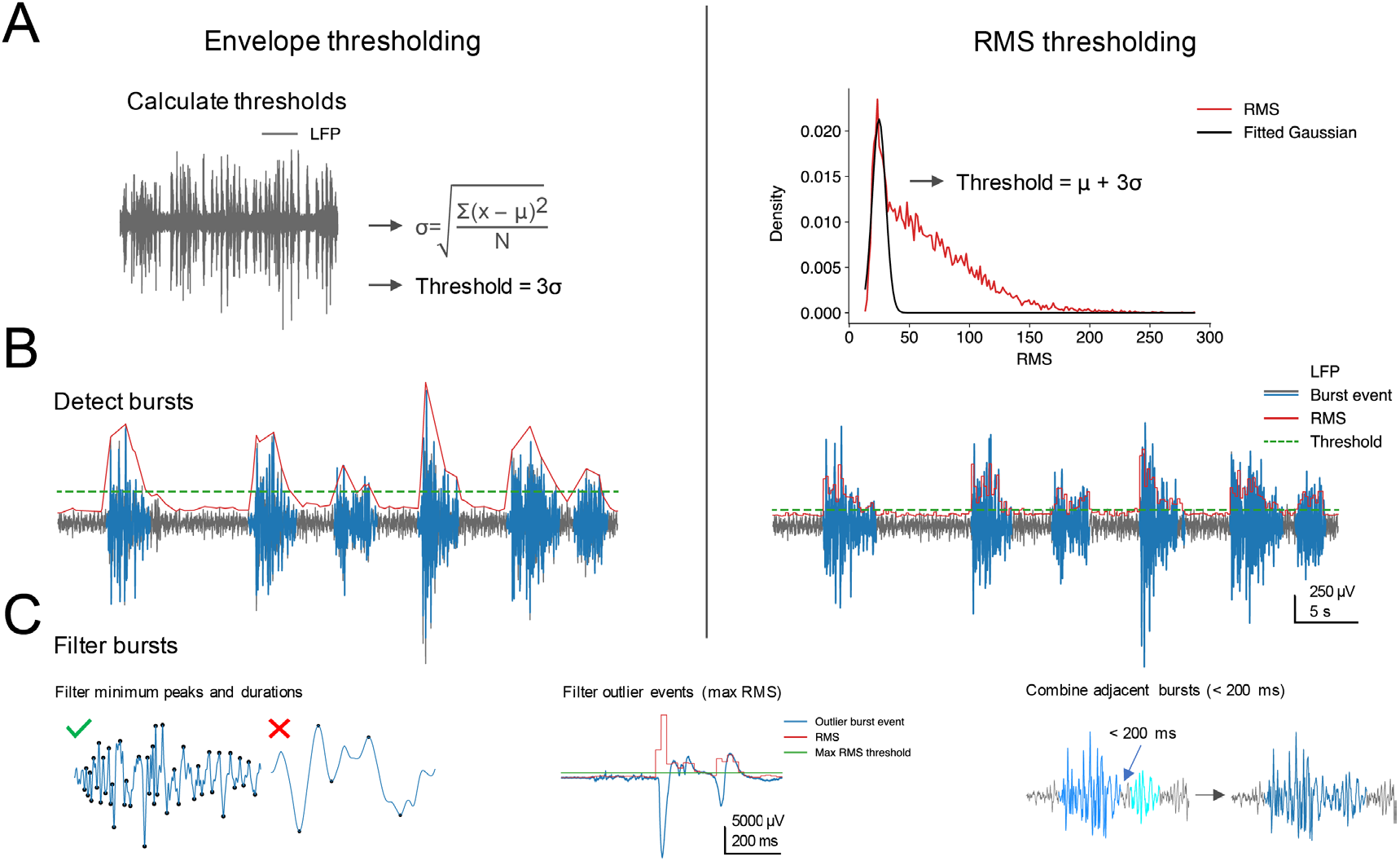
Burst detection algorithms used for analysis of recordings. (**A**) Two different algorithms were initially trialled. The bursting threshold was first computed either based on the standard deviation of the LFP signal (Envelope thresholding, left) or on the mean and standard deviation parameters of the fitted Gaussian (RMS thresholding, right). See also **Methods**. The RMS thresholding method was chosen for subsequent analyses as it is better validated in existing literature while being more parsimonious insofar as it does not require any per-recording parameters to be decided by the experimenter. (**B**) Burst events (in blue) were defined as those periods exceeding the threshold. (**C**) Putative burst events were subsequently checked to ensure they adhered to a minimum duration and number of peaks, while not exceeding the outlier RMS threshold. Finally, adjacent detected bursts found within 200 ms were deemed to be part of one burst event and combined.

**Supplemental Figure 2:**
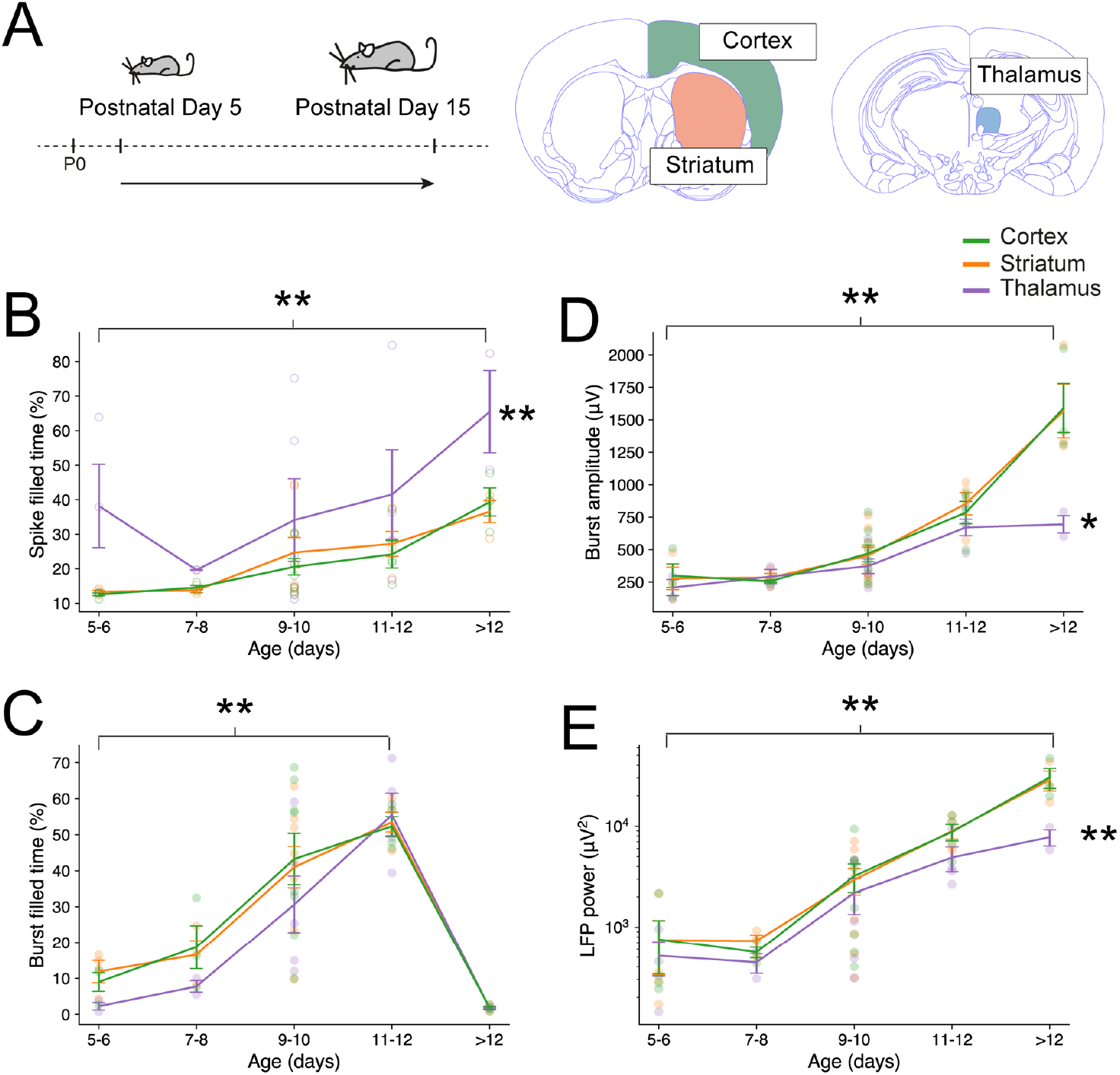
General increase in neural activity in all three brain regions across postnatal development. (**A**) Recordings were made from the cortex, dorsal striatum and intralaminar thalamus in mice between postnatal days (P)5-15. (**B**) Spike-filled time; fraction within 200 ms recording windows containing at least one spike event, increased significantly across postnatal development (F(4,45) = 4.66, p = 0.00312), while there was also a significant effect of group whereby spike-filled time was greater for the thalamus (F(2, 45) = 6.78, p = 0.00270). (**C**) Burst-filled time; proportion of time within each recording occupied by bursting events also increased significantly across postnatal development (F(4,42) = 26.1, p = 6.72e-11) before the main activity in all three brain regions transitioned to a continuous oscillatory pattern. (**D**) Mean burst amplitude, defined as the difference between the minimum and maximum peaks in the 4-100 Hz bandpass filtered LFP signals for each bursting event, increased significantly across postnatal development (F(4, 42) = 42.1, p = 3.61e-14). There was also a main effect of brain area (F(2, 42) = 4.88, p = 0.0125), where amplitude was lower across the thalamus, and a significant interaction effect of age and brain area (F(8, 42) = 2.50, p = 0.0258), likely driven by the disparity in amplitude at later developmental ages (> P12). (**E**) Local field potential (LFP) power in 4-100 Hz increased significantly across postnatal development (F(4,45) = 37.0, p = 1.06e-13). There was also a main effect of brain area (F(2, 45) = 4.21, p = 0.0212), where LFP power was lower for thalamus than for cortex or striatum, and a significant interaction effect of age and brain area (F(8, 45) = 3.07, p = 0.00754), where the increase in LFP power over time for thalamus was less pronounced as compared to cortex and striatum.

**Supplemental Figure 3:**
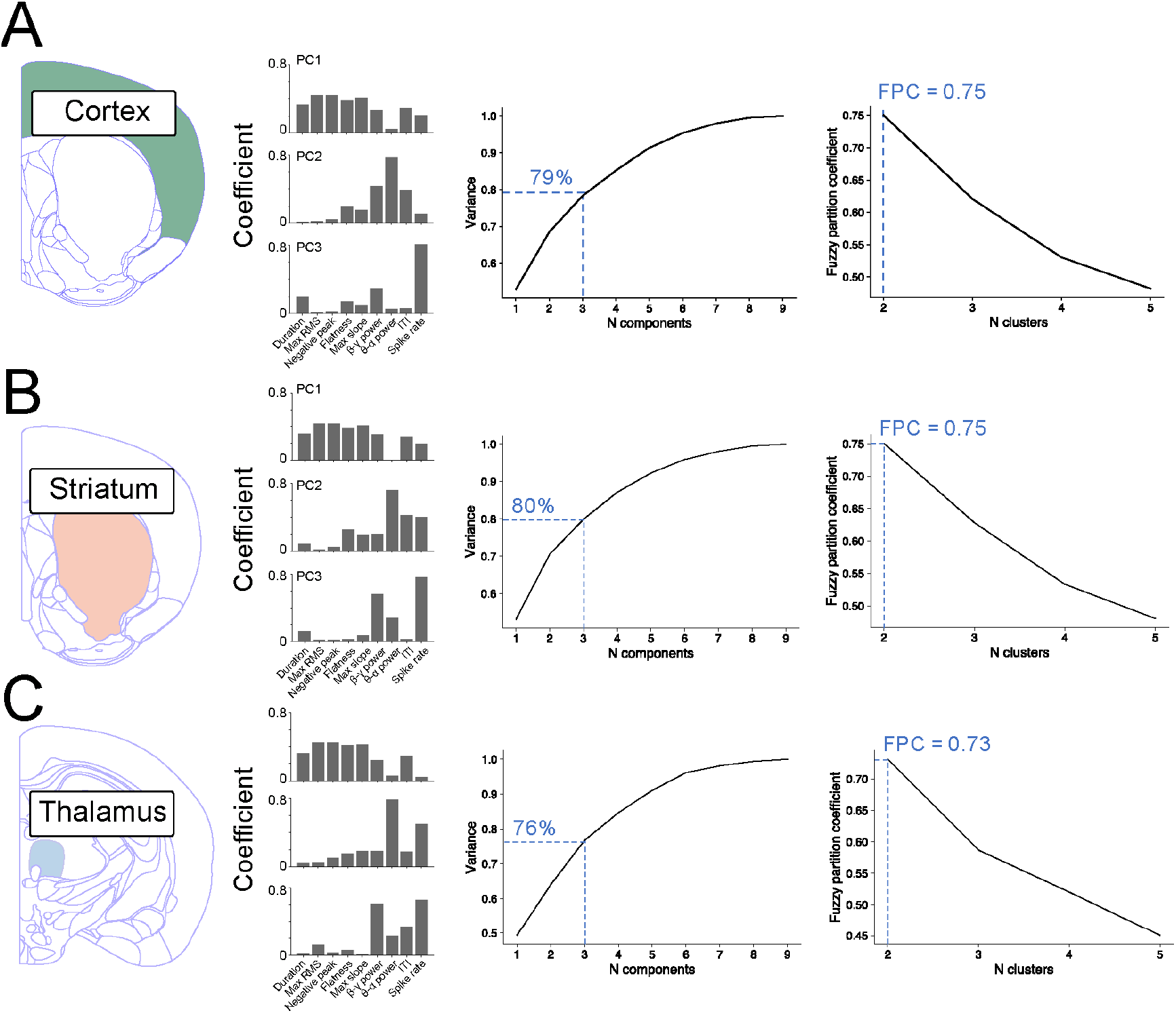
Principal component analysis (PCA) of detected events in cortex, striatum, and thalamus. Coefficient contribution of each feature to the first three principal components of detected (**A**) cortical, (**B**) striatal and (**C**) thalamic bursts and explained variance and fuzzy partition coefficients of detected bursts. Considering the contribution of different features to the first three principal components, PC1 appeared to represent a relatively uniform contribution across all features, albeit with minimal contribution of theta-alpha power. PC2 and PC3 were more dominated by specific features — namely theta-alpha power for PC2 and spike rate for PC3. Overall results were comparable across brain regions, though the explained variance for the first three components was slightly lower for the thalamic as compared with cortical and striatal burst events.

**Supplemental Figure 4:**
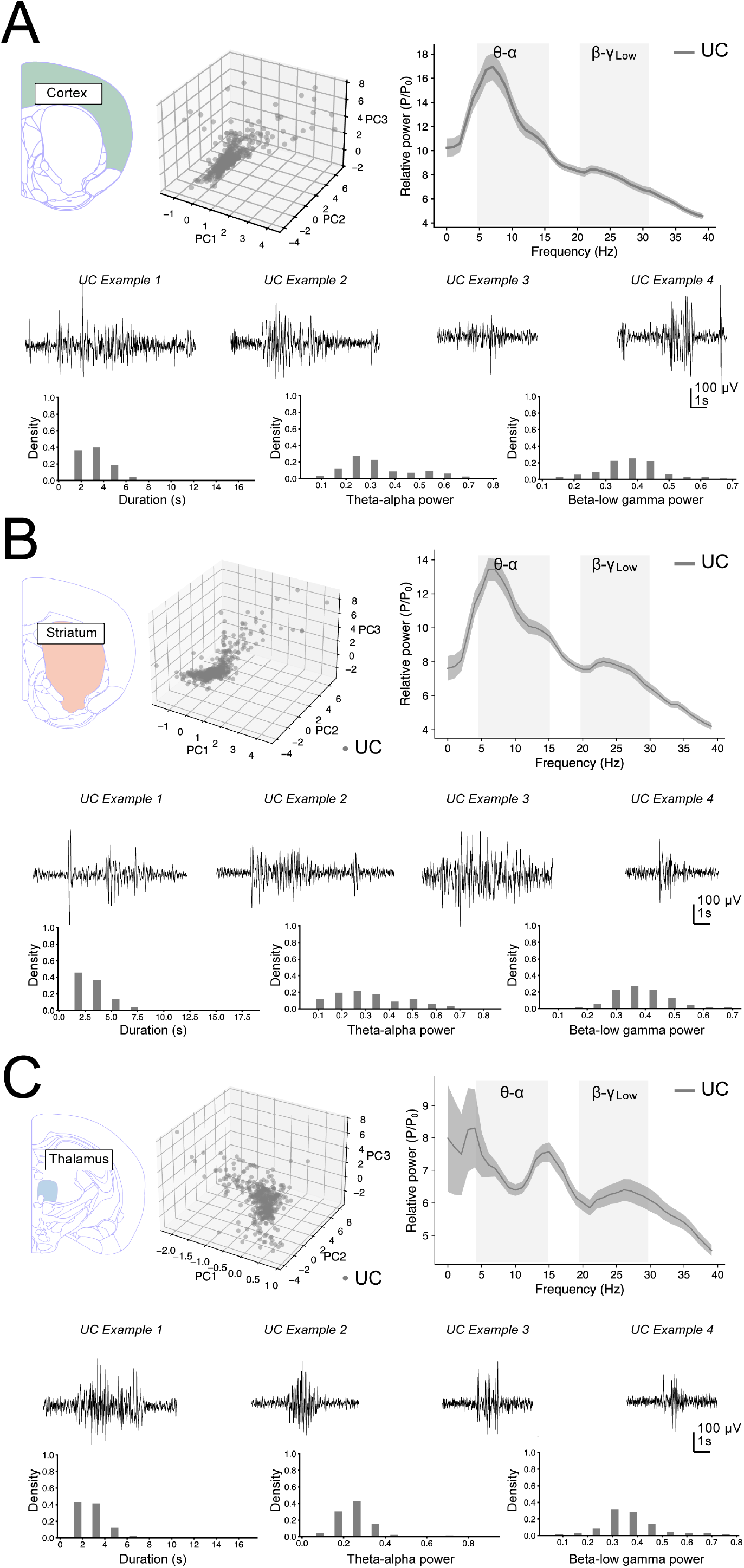
Characteristics of unclassified (UC) events. (**A**) Scatter plot of the first three principal components of striatal unclassified (UC) bursts (left). PSD of the mean normalized power across UC events (right). Note the peak at approximately theta-alpha frequency range (4-15 Hz). Four example UC events in striatum (middle). Histogram of the distributions of several key features across UC events including their duration, theta-alpha power, and beta-low gamma power (bottom). (**B**) Similar for UC events in striatum and (**C**) UC events in thalamus.

**Supplemental Figure 5:**
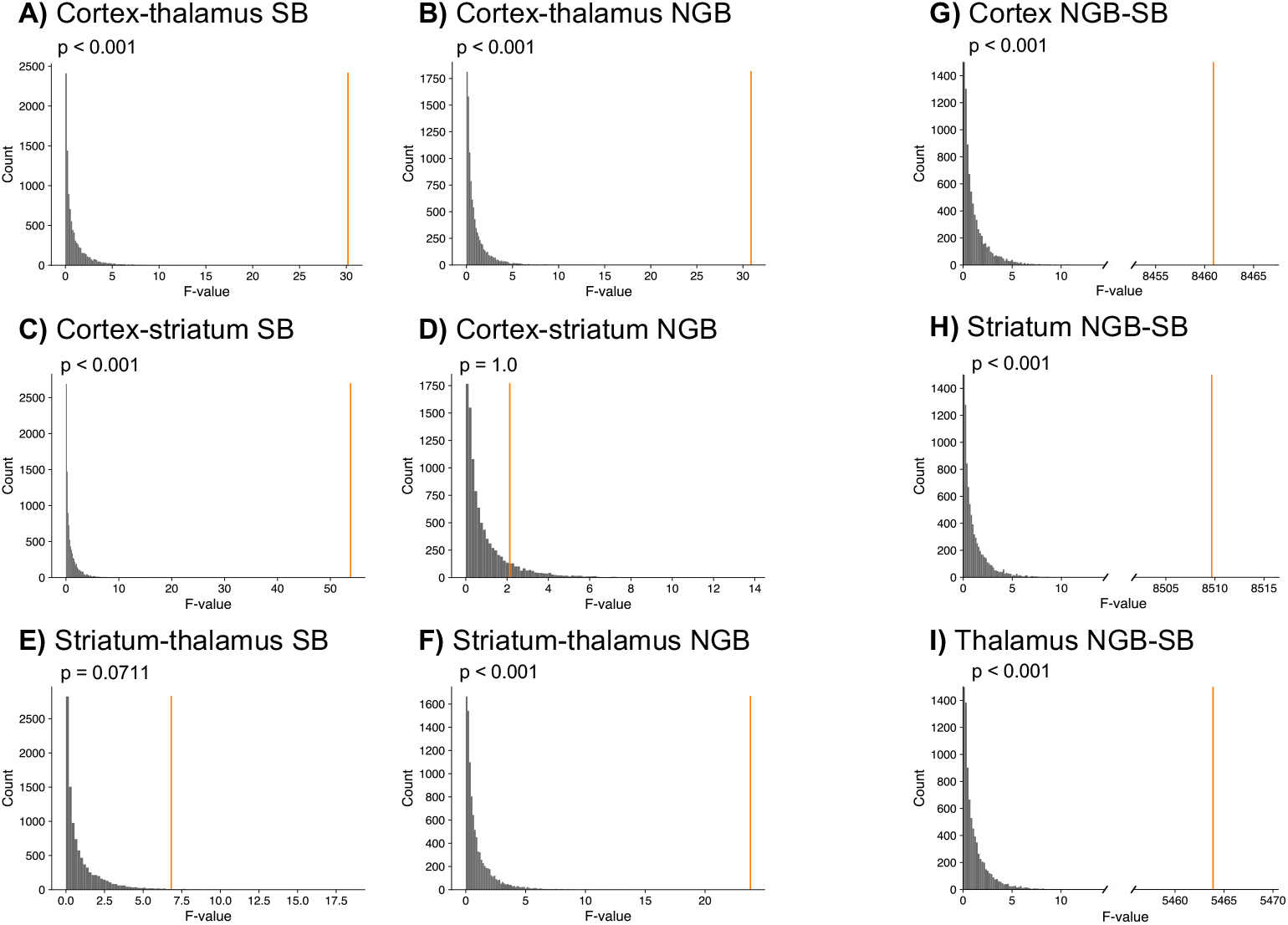
F-ratio comparison testing for NGB and SB events across striatum and thalamus. The significance was determined by shuffling the feature vectors between the two groups 10,000 and calculating the resulting F-statistic at each iteration to produce a null distribution. The significance threshold was defined as the 95^th^ percentile of this null distribution, with p-values given here corrected for multiple comparisons. There was a significant effect of brain area across all comparisons (**A, B, C, F**) except for NGB events between the cortex and striatum (**D**) as well as SB events between the striatum and thalamus (**E**) which did not reach significance after correction for multiple comparisons. F-ratio test for burst events across cortex (**G**), striatum (**H**) and thalamus (**I**). F-ratios are large for both cortex, striatum, and thalamus, indicating a robust effect of burst type.

**Supplemental Figure 6:**
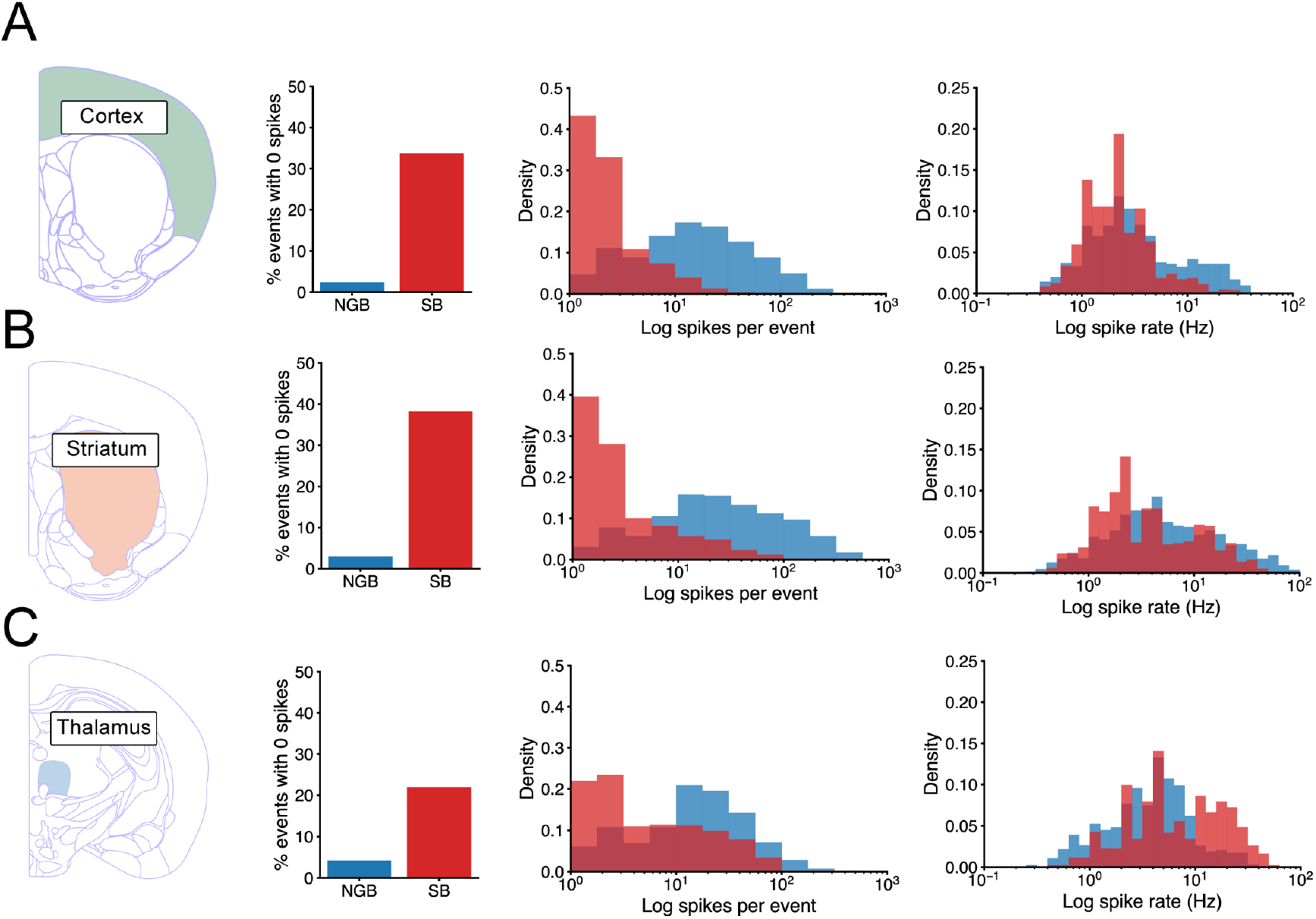
Characteristics of MUA activity. (**A**) For cortex the proportion of burst events that did not contain significant spiking (left). For those events that did contain spikes the distribution the number of spikes detected (middle). The distribution of spike frequencies in burst events (right). Same for (**B**) striatum and (**C**) thalamus.

**Supplemental Figure 7:**
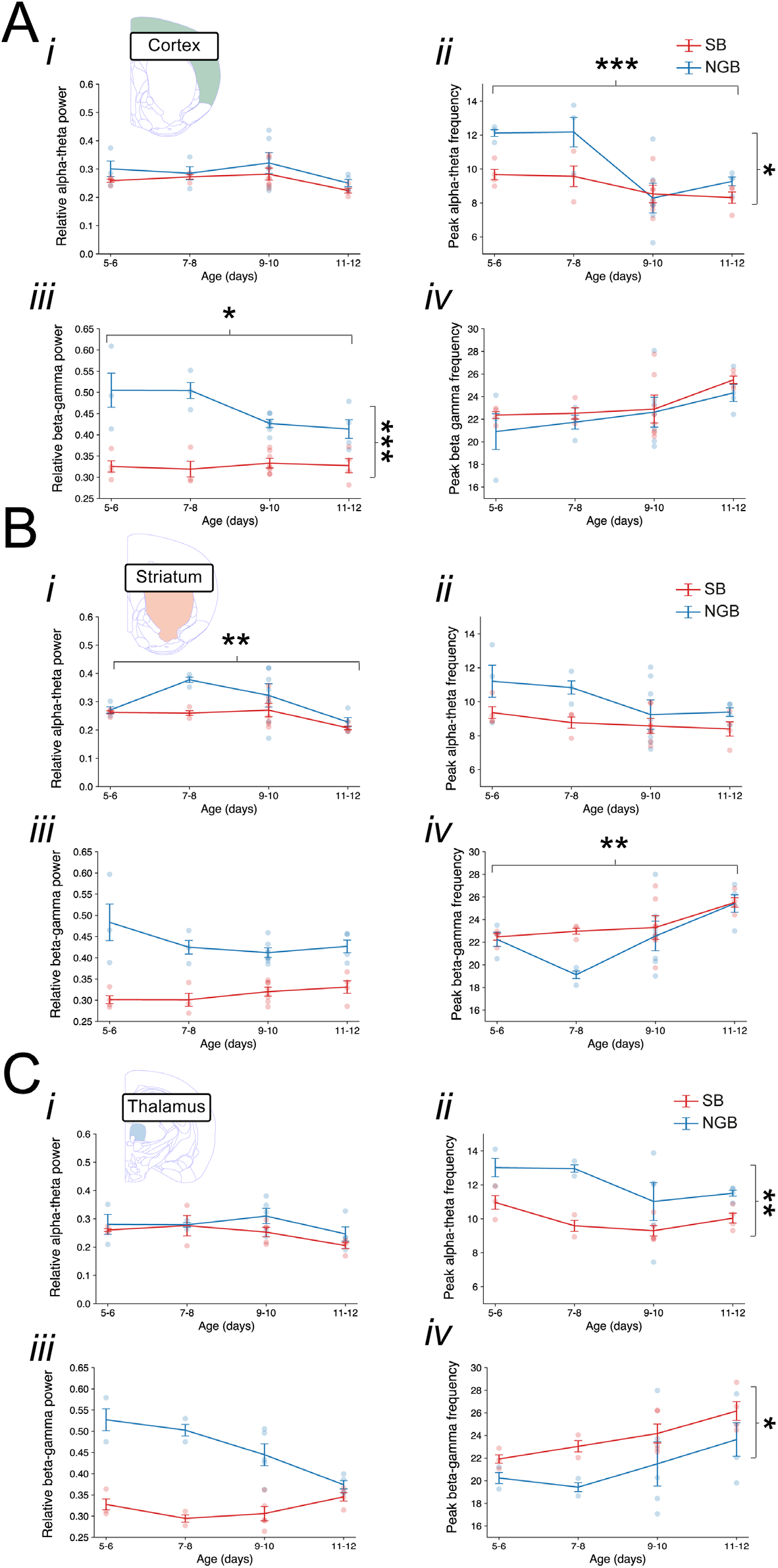
Developmental changes in burst properties. (**A**) Changes in the peak theta-alpha power (*i*) and frequency (*ii*) and peak beta-gamma power (*iii*) and frequency (*iv*) of those occurring in cortex. (**B**) Changes in the peak theta-alpha power (*i*) frequency (*ii*) and peak beta-low gamma power (*iii*) frequency (*iv*) of those occurring in striatum. The theta-alpha frequency component within striatal bursts across development exhibited a decrease in power (F(3,29) = 5.32, p = 0.00480) while the peak frequency did not change (F(3,29) = 2.21, p = 0.108). In contrast, the beta-low gamma frequency component within striatal bursts across development remained constant in power (F(3,29) = 2.34, p = 0.564), but significantly increased in frequency (F(3,29) = 6.56, p = 0.00160) increasing from 22.4 Hz to 25.5 Hz. (**C**) Changes in the peak theta-alpha power (*i*) frequency (*ii*) and peak beta-low gamma power (*iii*) frequency (*iv*) of those occurring in thalamus. Both the theta-alpha frequency and beta-low gamma frequency components within thalamic bursts remained constant in power (theta-alpha: F(3,19) = 1.86, p = 0.171 and beta-low gamma: F(3,19) = 2.54, p = 0.0752) and frequency (theta-alpha: F(3,19) = 2.20, p = 0.121 and beta-low gamma: F(3,19) = 2.69, p = 0.0752) across development. Note that the peak theta-alpha frequency was greater for NGB over SB events (F(1, 19) = 14.2, p = 0.00131) and the peak beta-low gamma frequency slightly faster for SB than NGB events (F(1, 19 = 5.93, p = 0.0249), though the overall power in these SB events was lower (F(1, 19) = 64.8, p = 1.54e-07). Interestingly, NGB beta-low gamma power appeared to decrease while remaining constant for SB events over time (F(3, 19) = 6.52, p = 0.00323).

**Supplemental Figure 8:**
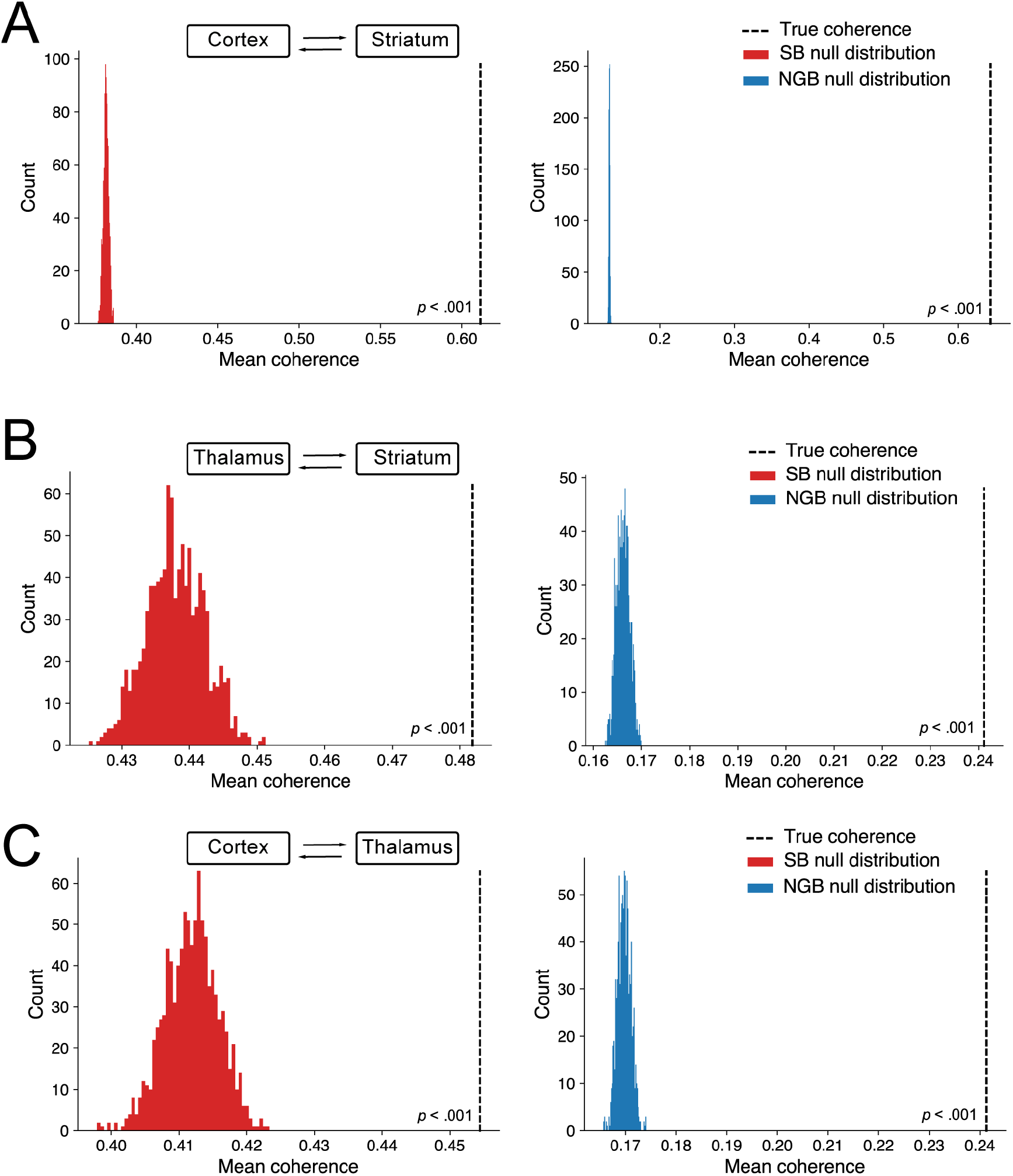
Permutation test for mean coherence across SB and NGB events. (**A**) Pairs of cortical-striatal events. (**B**) Pairs of thalamic-striatal events. (**C**) Pairs of cortical-thalamic events. Pairs of NGB and SB events were randomly shuffled 1000 times, with the mean coherence across all shuffled pairs computed on each iteration to produce a null distribution. The significance threshold was defined as the 95^th^ percentile of this null distribution.

**Supplemental Figure 9:**
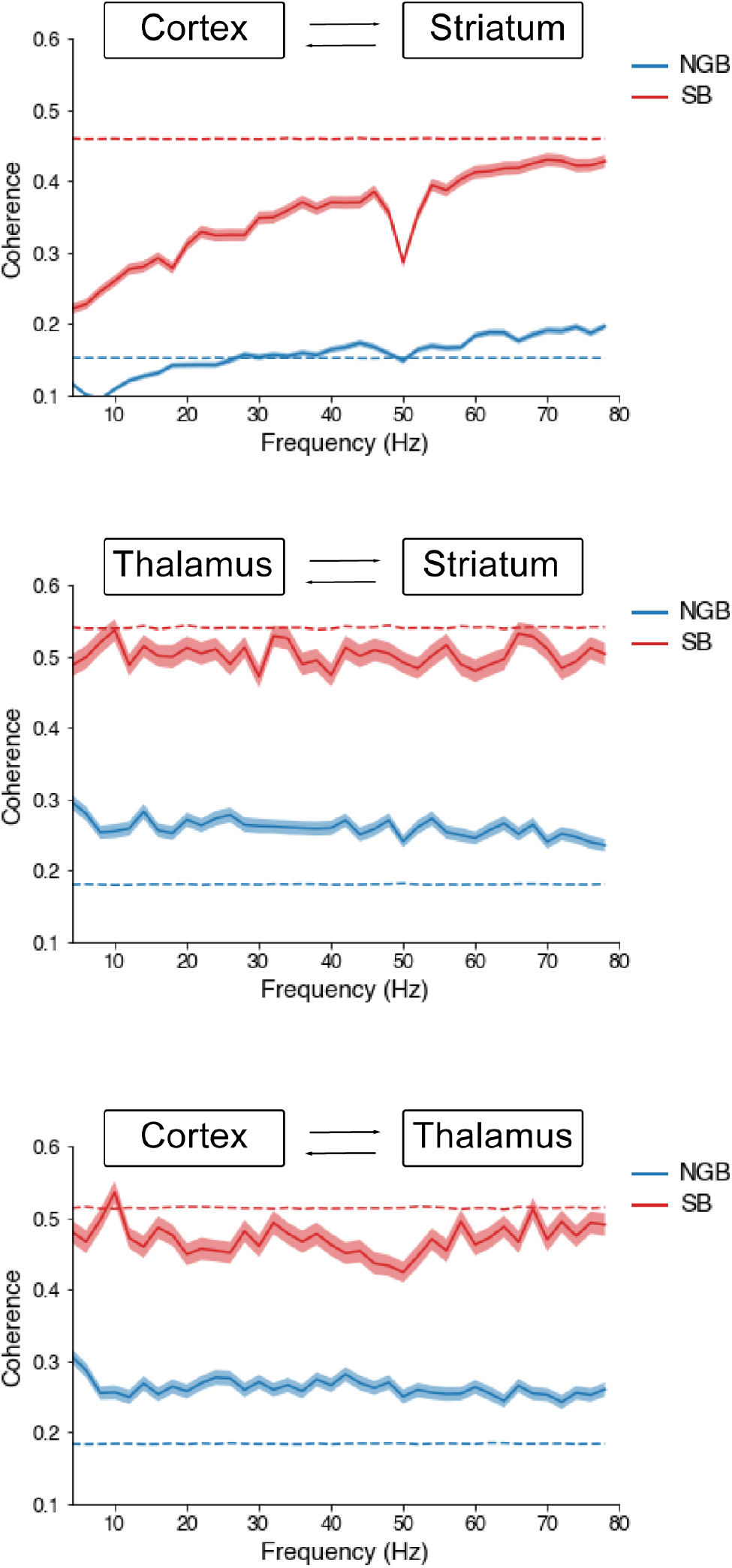
Imaginary cross-spectral coherence for SB and NGB events. Imaginary coherence analysis was performed on co-occurring burst events, segregated into NGB-NGB and SB-SB events to reveal potential non-zero lag interactions. Shuffled data (dashed line) signifies the 95^th^ percentile of the shuffled (null) distribution. Overall, NGB events appear to be significantly coherent while SB events do not exceed coherence values that would be expected by chance assuming no coherence.

**Supplemental Figure 10:**
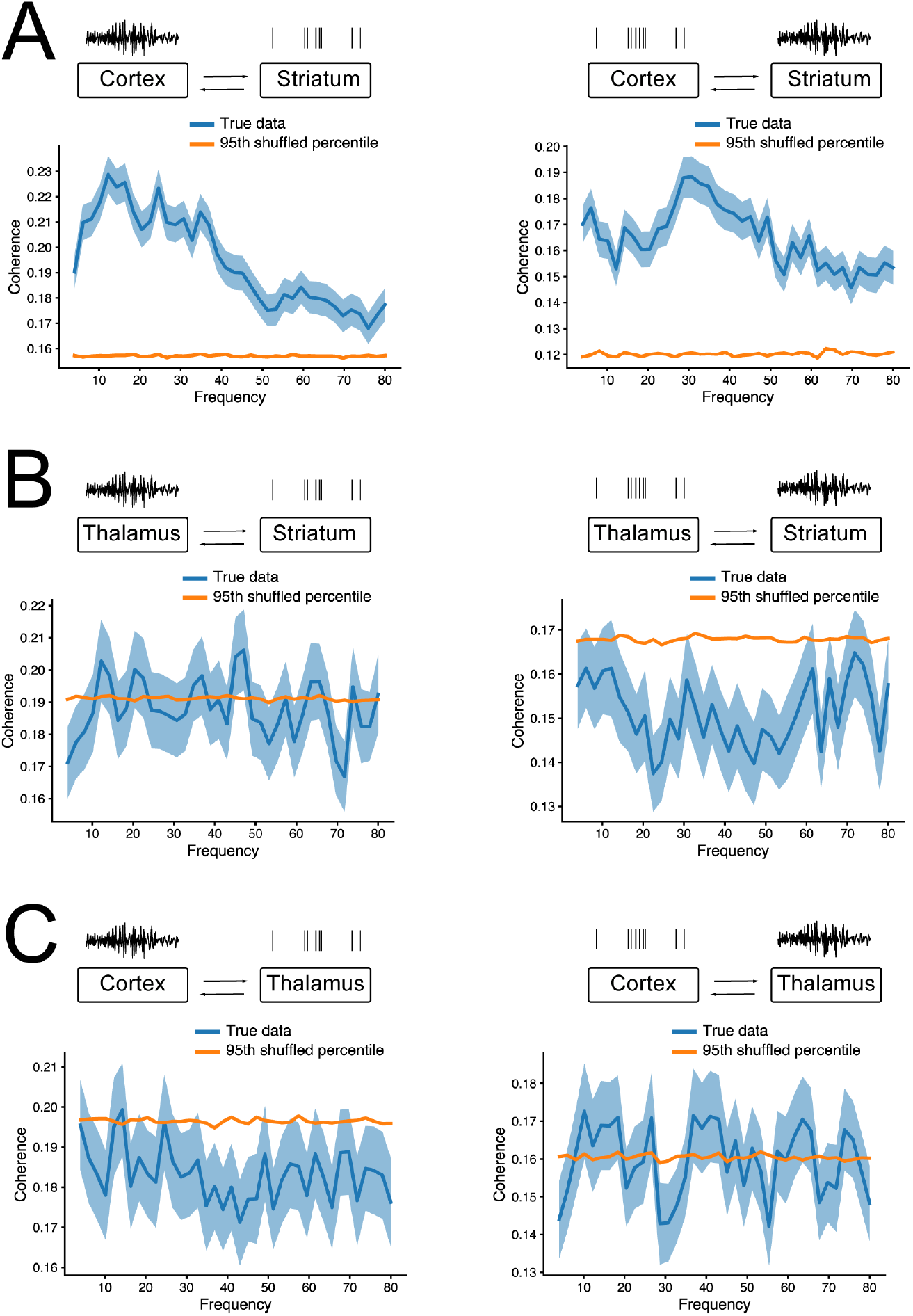
Spike-field coherence analysis suggest interactions between brain regions. (**A**) Spike-field coherence for cortico-striatal interactions. Shuffled lines indicate 95^th^ percentile range based on null distribution (shuffled data). Note the significant interaction of the cortex LFP with spiking in the striatum (left, p<0.001) and the significant interaction of the cortical spiking with striatal LFP (right, p<0.001). (**B**) Spike-field coherence for striatal-thalamic interactions. No significant interaction was observed for thalamic LFP with spiking in the striatum (left, p<0.431) or thalamic spiking with striatal LFP (right, p<0.084). (**C**) Spike-field coherence for cortico-thalamic interactions. Spike-field coherence for cortico-striatal interactions. Although no significant interaction was observed for cortex LFP with spiking in the thalamus (left, p<0.431) we find a significant interaction of the cortical spiking with thalamic LFP (right, p<0.001).

**Supplemental Figure 11:**
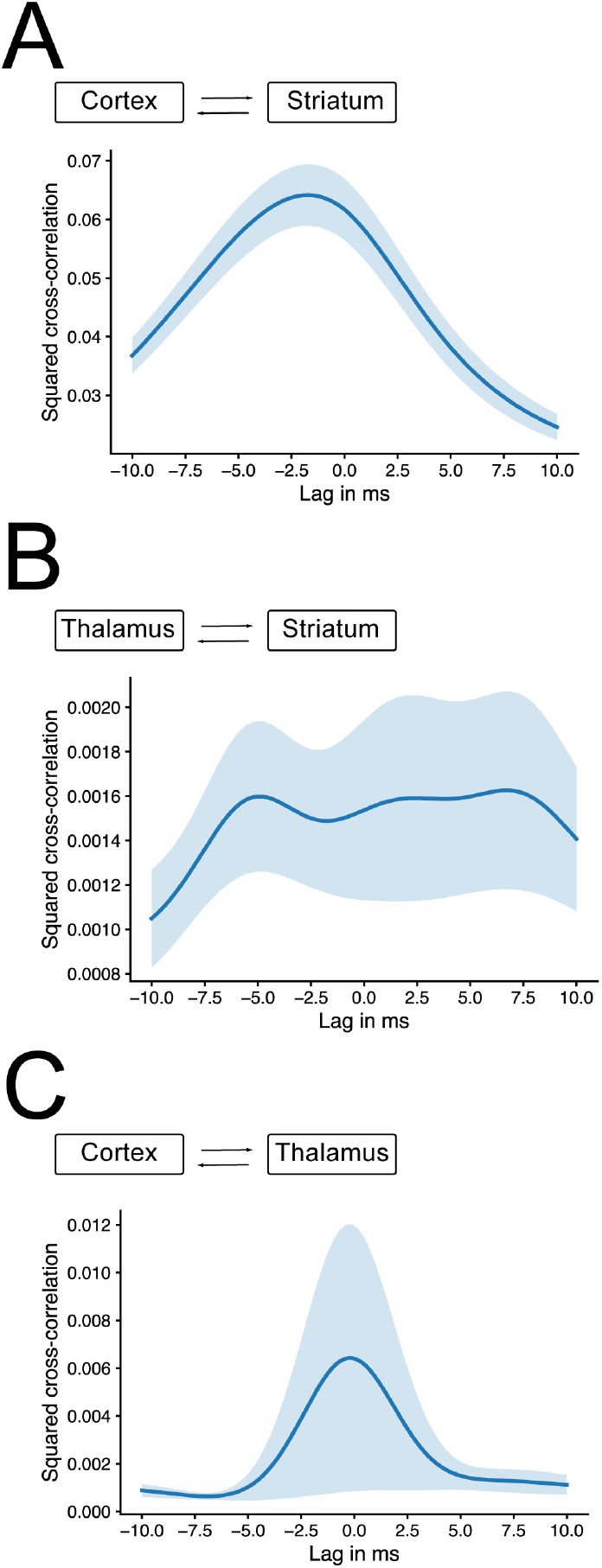
Spike-train cross-correlation analysis suggests strong impact of cortical on striatal activity. (**A**) Spike train cross-correlation (convolved with a Gaussian kernel) between cortex and striatum suggest a strong drive of cortex on striatal activity (peak of mean = −1.75 ms, one-sample *t*-test, p = 4.26e-06). (**B**) Spike train cross-correlation between thalamus and striatum does not reveal evidence for one brain region driving other more strongly (one-sample *t*-test, p = 0.80). (**C**) Spike train cross-correlation between cortex and thalamus does not reveal evidence for one brain region driving other more strongly (peak of mean = −0.22 ms, one-sample *t*-test, p = 0.49).

### Supplemental Tables

**Supplemental Table 1:**
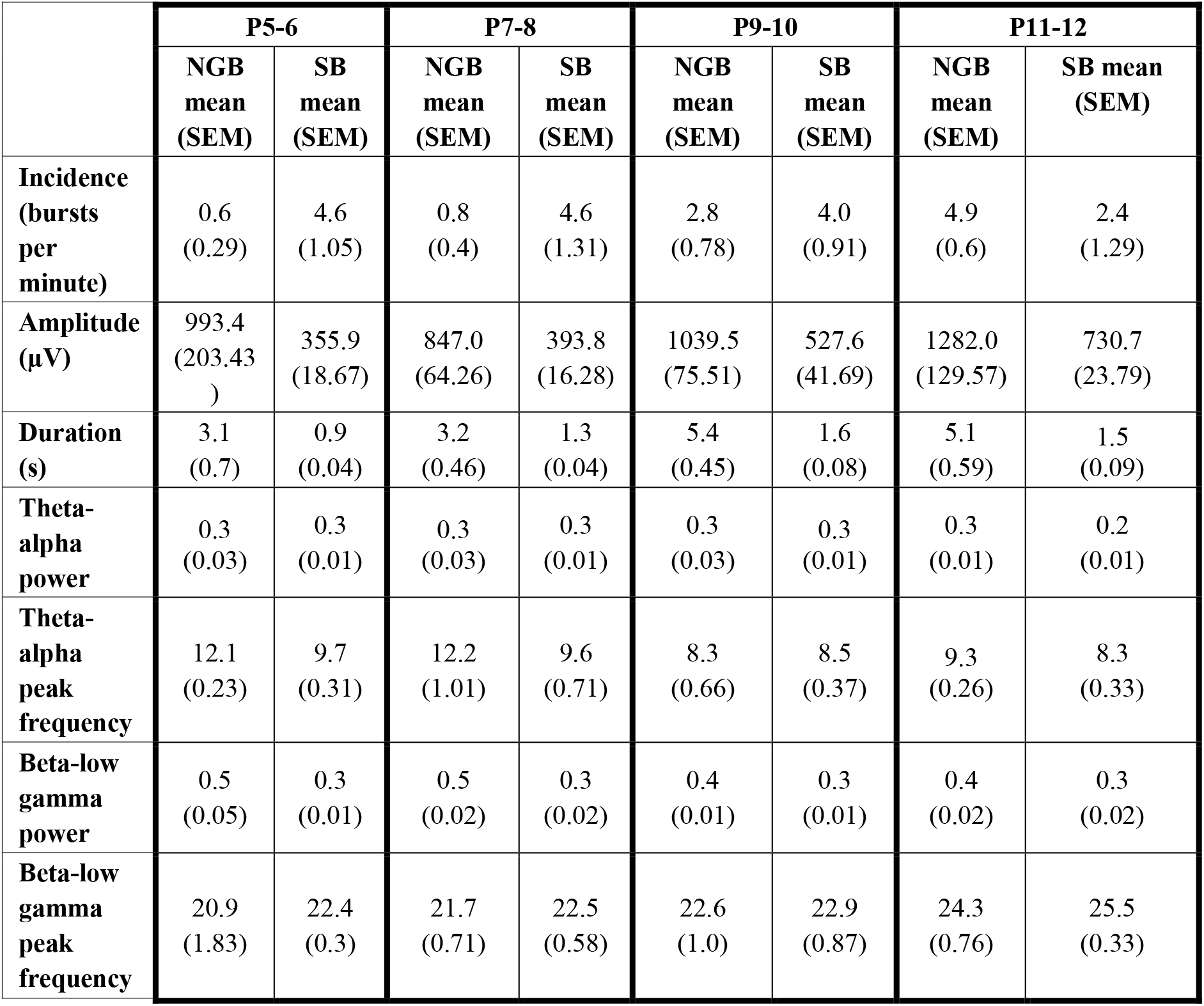
Parameters for cortical burst events.

|  | P5-6 |  | P7-8 |  | P9-10 |  | P11-12 |  |
| --- | --- | --- | --- | --- | --- | --- | --- | --- |
|  | NGB mean (SEM) | SB mean (SEM) | NGB mean (SEM) | SB mean (SEM) | NGB mean (SEM) | SB mean (SEM) | NGB mean (SEM) | SB mean (SEM) |
| **Incidence (bursts per minute)** | 0.6 (0.29) | 4.6 (1.05) | 0.8 (0.4) | 4.6 (1.31) | 2.8 (0.78) | 4.0 (0.91) | 4.9 (0.6) | 2.4 (1.29) |
| **Amplitude (μV)** | 993.4 (203.43) | 355.9 (18.67) | 847.0 (64.26) | 393.8 (16.28) | 1039.5 (75.51) | 527.6 (41.69) | 1282.0 (129.57) | 730.7 (23.79) |
| **Duration (s)** | 3.1 (0.7) | 0.9 (0.04) | 3.2 (0.46) | 1.3 (0.04) | 5.4 (0.45) | 1.6 (0.08) | 5.1 (0.59) | 1.5 (0.09) |
| **Theta-alpha power** | 0.3 (0.03) | 0.3 (0.01) | 0.3 (0.03) | 0.3 (0.01) | 0.3 (0.03) | 0.3 (0.01) | 0.3 (0.01) | 0.2 (0.01) |
| **Theta-alpha peak frequency** | 12.1 (0.23) | 9.7 (0.31) | 12.2 (1.01) | 9.6 (0.71) | 8.3 (0.66) | 8.5 (0.37) | 9.3 (0.26) | 8.3 (0.33) |
| **Beta-low gamma power** | 0.5 (0.05) | 0.3 (0.01) | 0.5 (0.02) | 0.3 (0.02) | 0.4 (0.01) | 0.3 (0.01) | 0.4 (0.02) | 0.3 (0.02) |
| **Beta-low gamma peak frequency** | 20.9 (1.83) | 22.4 (0.3) | 21.7 (0.71) | 22.5 (0.58) | 22.6 (1.0) | 22.9 (0.87) | 24.3 (0.76) | 25.5 (0.33) |

**Supplemental Table 2:** Parameters for striatal burst events.

|  | P5-6 |  | P7-8 |  | P9-10 |  | P11-12 |  |
| --- | --- | --- | --- | --- | --- | --- | --- | --- |
|  | NGB<br>mean<br>(SEM) | SB<br>mean<br>(SEM) | NGB<br>mean<br>(SEM) | SB<br>mean<br>(SEM) | NGB<br>mean<br>(SEM) | SB<br>mean<br>(SEM) | NGB<br>mean<br>(SEM) | SB<br>mean<br>(SEM) |
| **Incidence**<br>**(bursts**<br>**per**<br>**minute)** | 0.7<br>(0.35) | 5.6<br>(0.99) | 0.8<br>(0.23) | 4.3<br>(1.01) | 0.7<br>(0.35) | 5.6<br>(0.99) | 0.8<br>(0.23) | 4.3<br>(1.01) |
| **Amplitude**<br>**(μV)** | 1031.5<br>(211.15) | 343.9<br>(18.91) | 946.8<br>(47.64) | 435.1<br>(23.38) | 1031.5<br>(211.15) | 343.9<br>(18.91) | 946.8<br>(47.64) | 435.1<br>(23.38) |
| **Duration**<br>**(s)** | 2.5<br>(0.17) | 1.0<br>(0.04) | 4.0<br>(0.07) | 1.3<br>(0.05) | 2.5<br>(0.17) | 1.0<br>(0.04) | 4.0<br>(0.07) | 1.3<br>(0.05) |
| **Theta-**<br>**alpha**<br>**power** | 0.3<br>(0.01) | 0.3<br>(0.01) | 0.4<br>(0.01) | 0.3<br>(0.01) | 0.3<br>(0.01) | 0.3<br>(0.01) | 0.4<br>(0.01) | 0.3<br>(0.01) |
| **Theta-**<br>**alpha**<br>**peak**<br>**frequency** | 11.2<br>(1.09) | 9.4<br>(0.35) | 10.8<br>(0.46) | 8.8<br>(0.38) | 11.2<br>(1.09) | 9.4<br>(0.35) | 10.8<br>(0.46) | 8.8<br>(0.38) |
| **Beta-low**<br>**gamma**<br>**power** | 0.5<br>(0.05) | 0.3<br>(0.01) | 0.4<br>(0.02) | 0.3<br>(0.02) | 0.5<br>(0.05) | 0.3<br>(0.01) | 0.4<br>(0.02) | 0.3<br>(0.02) |
| **Beta-low**<br>**gamma**<br>**peak**<br>**frequency** | 22.3<br>(0.73) | 22.5<br>(0.28) | 19.1<br>(0.39) | 23.0<br>(0.31) | 22.3<br>(0.73) | 22.5<br>(0.28) | 19.1<br>(0.39) | 23.0<br>(0.31) |

**Supplemental Table 3:** Parameters for thalamic burst events.

|  | P5-6 |  | P7-8 |  | P9-10 |  | P11-12 |  |
| --- | --- | --- | --- | --- | --- | --- | --- | --- |
|  | NGB<br>mean<br>(SEM) | SB<br>mean<br>(SEM) | NGB<br>mean<br>(SEM) | SB<br>mean<br>(SEM) | NGB<br>mean<br>(SEM) | SB<br>mean<br>(SEM) | NGB<br>mean<br>(SEM) | SB<br>mean<br>(SEM) |
| **Incidence**<br>**(bursts**<br>**per**<br>**minute)** | 0.4<br>(0.25) | 2.2<br>(0.97) | 0.4<br>(0.02) | 1.8<br>(0.7) | 2.3<br>(0.75) | 4.0<br>(0.48) | 5.4<br>(1.06) | 2.3<br>(1.01) |
| **Amplitude**<br>**(μV)** | 957.1<br>(168.16) | 452.6<br>(44.98) | 920.4<br>(139.7) | 412.3<br>(44.56) | 987.2<br>(57.47) | 489.4<br>(77.99) | 1043.2<br>(34.44) | 617.1<br>(46.85) |
| **Duration**<br>**(s)** | 1.2<br>(0.07) | 0.7<br>(0.12) | 2.9<br>(0.19) | 1.1<br>(0.07) | 4.5<br>(1.17) | 1.4<br>(0.1) | 4.6<br>(0.28) | 1.5<br>(0.09) |
| **Theta-**<br>**alpha**<br>**power** | 0.3<br>(0.05) | 0.3<br>(0.01) | 0.3<br>(0.01) | 0.3<br>(0.05) | 0.3<br>(0.02) | 0.3<br>(0.01) | 0.2<br>(0.03) | 0.2<br>(0.01) |
| **Theta-**<br>**alpha**<br>**peak**<br>**frequency** | 13.0<br>(0.77) | 11.0<br>(0.46) | 13.0<br>(0.31) | 9.6<br>(0.46) | 11.0<br>(1.0) | 9.3<br>(0.28) | 11.5<br>(0.18) | 10.0<br>(0.29) |
| **Beta-low**<br>**gamma**<br>**power** | 0.5<br>(0.04) | 0.3<br>(0.01) | 0.5<br>(0.02) | 0.3<br>(0.01) | 0.4<br>(0.02) | 0.3<br>(0.02) | 0.4<br>(0.01) | 0.3<br>(0.01) |
| **Beta-low**<br>**gamma**<br>**peak**<br>**frequency** | 20.2<br>(0.69) | 21.9<br>(0.43) | 19.4<br>(0.55) | 23.0<br>(0.71) | 21.5<br>(1.76) | 24.2<br>(0.74) | 23.6<br>(1.49) | 26.2<br>(0.83) |

**Supplemental Table 4:** Statistics for cortical burst events.

|  | **NGB mean**<br>**(SEM)** | **SB mean**<br>**(SEM)** | **df** | **Welch's**<br>**t** | **p**<br>**(uncorrected)** | **Cohen's d** |
| --- | --- | --- | --- | --- | --- | --- |
| **Duration**<br>**(s)** | 5.52<br>(0.0844) | 1.44 (0.0192) | 1.83e+03 | 47.2 | 9.08e-319 | 1.63 |
| **Max RMS**<br>**(μV)** | 193 (1.48) | 56.1 (0.628) | 2.25e+03 | 85.3 | 0 | 2.89 |
| **Negative**<br>**peak (μV)** | -420 (3.19) | -126 (1.33) | 2.23e+03 | -85.4 | 0 | -2.89 |
| **Flatness** | 0.213<br>(0.00203) | 0.538<br>(0.00398) | 3.25e+03 | -72.8 | 0 | -2.24 |
| **Max slope** | 130 (1.23) | 42.8 (0.377) | 1.97e+03 | 68 | 0 | 2.33 |
| **Beta/low-**<br>**gamma**<br>**power** | 0.428<br>(0.00164) | 0.324<br>(0.00181) | 3.88e+03 | 42.5 | 9.88e-324 | 1.35 |
| **Theta-**<br>**alpha**<br>**power** | 0.273<br>(0.00179) | 0.268<br>(0.00188) | 3.85e+03 | 1.92 | 0.0549 | 0.0613 |
| **Inter-**<br>**trough-**<br>**interval (s)** | 0.0662<br>(0.000181) | 0.0551<br>(0.000177) | 3.78e+03 | 43.9 | 0 | 1.41 |
| **Spikes s<sup>-1</sup>** | 5.21<br>(0.166) | 1.35 (0.0508) | 1.97e+03 | 22.2 | 8.58e-98 | 0.761 |

**Supplemental Table 5:** Statistics for striatal burst events.

|  | **NGB mean**<br>**(SEM)** | **SB mean**<br>**(SEM)** | **df** | **Welch's**<br>**t** | **p**<br>**(uncorrected)** | **Cohen's**<br>**d** |
| --- | --- | --- | --- | --- | --- | --- |
| **Duration (s)** | 5.61<br>(0.0907) | 1.31<br>(0.0189) | 1.77e+03 | 46.4 | 4.64e-308 | 1.61 |
| **Maximum**<br>**RMS (μV)** | 190<br>(1.52) | 49.3<br>(0.56) | 2.08e+03 | 87.2 | 0 | 2.99 |
| **Negative peak**<br>**(μV)** | -417<br>(3.4) | -112<br>(1.21) | 2.05e+03 | -84.4 | 0 | -2.9 |
| **Flatness** | 0.213<br>(0.00207) | 0.576<br>(0.00401) | 3.27e+03 | -80.3 | 0 | -2.47 |
| **Maximum**<br>**slope** | 136<br>(1.42) | 41.8<br>(0.361) | 1.84e+03 | 64.1 | 0 | 2.22 |
| **Relative beta-**<br>**low gamma**<br>**power** | 0.424<br>(0.00159) | 0.306<br>(0.00171) | 3.85e+03 | 50.5 | 0 | 1.61 |
| **Relative**<br>**theta-alpha**<br>**power** | 0.268<br>(0.00227) | 0.264<br>(0.00193) | 3.51e+03 | 1.34 | 0.179 | 0.0438 |
| **Inter-trough-**<br>**interval (s)** | 0.0643<br>(0.000179) | 0.0527<br>(0.000162) | 3.63e+03 | 48.4 | 0 | 1.57 |
| **Spikes s<sup>-1</sup>** | 9.75<br>(0.341) | 2.51<br>(0.116) | 2.01e+03 | 20.1 | 6.14e-82 | 0.69 |

**Supplemental Table 6:** Statistics for thalamic burst events.

|  | **NGB mean**<br>**(SEM)** | **SB mean**<br>**(SEM)** | **df** | **Welch's**<br>**t** | **p**<br>**(uncorrected)** | **Cohen's**<br>**d** |
| --- | --- | --- | --- | --- | --- | --- |
| **Duration**<br>**(s)** | 4.85<br>(0.0835) | 1.29<br>(0.0233) | 1.54e+03 | 41 | 2.89e-249 | 1.59 |
| **Maximum**<br>**RMS (μV)** | 165<br>(1.12) | 50.4<br>(0.71) | 2.27e+03 | 86.9 | 0 | 3.35 |
| **Negative**<br>**peak (μV)** | -406<br>(3.7) | -115<br>(1.52) | 1.78e+03 | -72.8 | 0 | -2.81 |
| **Flatness** | 0.224<br>(0.00234) | 0.612<br>(0.00474) | 2.01e+03 | -73.5 | 0 | -2.81 |
| **Maximum**<br>**slope** | 131<br>(1.3) | 44.7<br>(0.504) | 1.73e+03 | 61.6 | 0 | 2.38 |
| **Relative**<br>**beta-low**<br>**gamma**<br>**power** | 0.396<br>(0.00194) | 0.315<br>(0.00227) | 2.67e+03 | 27.2 | 1.47e-143 | 1.04 |
| **Relative**<br>**theta-alpha**<br>**power** | 0.259<br>(0.0023) | 0.24<br>(0.00236) | 2.72e+03 | 5.48 | 4.61e-08 | 0.21 |
| **Inter-**<br>**trough-**<br>**interval (s)** | 0.0591<br>(0.000169) | 0.0505<br>(0.000183) | 2.71e+03 | 34.6 | 3.36e-217 | 1.33 |
| **Spikes $s^{-1}$** | 4.86<br>(0.146) | 6.51<br>(0.243) | 2.26e+03 | -5.81 | 7.09e-09 | -0.222 |

**Supplemental Table 7:** Statistics for striatum versus thalamic NGB events.

|  | **Striatum**<br>**mean (SEM)** | **Thalamus**<br>**mean (SEM)** | **df** | **Welch's**<br>**t** | **p**<br>**(uncorrected)** | **Cohen's d** |
| --- | --- | --- | --- | --- | --- | --- |
| **Duration**<br>**(s)** | 5.61 (0.0907) | 4.85 (0.0835) | 2.97e+03 | 6.17 | 7.75e-10 | 0.226 |
| **Maximum**<br>**RMS (μV)** | 190 (1.52) | 165 (1.12) | 2.85e+03 | 13.2 | 7.37e-39 | 0.479 |
| **Negative**<br>**peak (μV)** | -417 (3.4) | -406 (3.7) | 2.87e+03 | -2.15 | 0.0313 | -0.0794 |
| **Flatness** | 0.213<br>(0.00207) | 0.224<br>(0.00234) | 2.82e+03 | -3.3 | 0.000983 | -0.122 |
| **Maximum**<br>**slope** | 136 (1.42) | 131 (1.3) | 2.97e+03 | 2.61 | 0.00908 | 0.0955 |
| **Relative**<br>**beta-low**<br>**gamma**<br>**power** | 0.424<br>(0.00159) | 0.396<br>(0.00194) | 2.72e+03 | 11.3 | 8.37e-29 | 0.418 |
| **Relative**<br>**theta-**<br>**alpha**<br>**power** | 0.268<br>(0.00227) | 0.259 (0.0023) | 2.93e+03 | 3.01 | 0.00263 | 0.111 |
| **Inter-**<br>**trough-**<br>**interval**<br>**(s)** | 0.0643<br>(0.000179) | 0.0591<br>(0.000169) | 2.96e+03 | 21.2 | 2.93e-93 | 0.778 |
| **Spikes s<sup>-1</sup>** | 9.75 (0.341) | 4.86 (0.146) | 2.19e+03 | 13.2 | 4.06e-38 | 0.467 |

**Supplemental Table 8:** Statistics for striatum versus thalamic SB events.

|  | **Striatum**<br>**mean (SEM)** | **Thalamus**<br>**mean (SEM)** | **df** | **Welch's**<br>**t** | **p**<br>**(uncorrected)** | **Cohen's**<br>**d** |
| --- | --- | --- | --- | --- | --- | --- |
| **Duration (s)** | 1.31 (0.0189) | 1.29 (0.0233) | 3e+03 | 0.736 | 0.462 | 0.0251 |
| **Max RMS**<br>**(μV)** | 49.3 (0.56) | 50.4 (0.71) | 2.94e+03 | -1.21 | 0.225 | -0.0415 |
| **Negative**<br>**peak (μV)** | -112 (1.21) | -115 (1.52) | 2.96e+03 | 1.53 | 0.125 | 0.0524 |
| **Flatness** | 0.576<br>(0.00401) | 0.612<br>(0.00474) | 3.1e+03 | -5.8 | 7.5e-09 | -0.196 |
| **Max slope** | 41.8 (0.361) | 44.7 (0.504) | 2.73e+03 | -4.67 | 3.18e-06 | -0.161 |
| **Beta/low-**<br>**gamma**<br>**power** | 0.306<br>(0.00171) | 0.315<br>(0.00227) | 2.84e+03 | -3.05 | 0.00234 | -0.105 |
| **Theta-alpha**<br>**power** | 0.264<br>(0.00193) | 0.24<br>(0.00236) | 3.03e+03 | 7.8 | 8.3e-15 | 0.265 |
| **Inter-**<br>**trough-**<br>**interval (s)** | 0.0527<br>(0.000162) | 0.0505<br>(0.000183) | 3.19e+03 | 8.85 | 1.4e-18 | 0.298 |
| **Spikes $s^{-1}$** | 2.51 (0.116) | 6.51 (0.243) | 2.02e+03 | -14.9 | 1.41e-47 | -0.536 |

**Supplemental Table 9:**
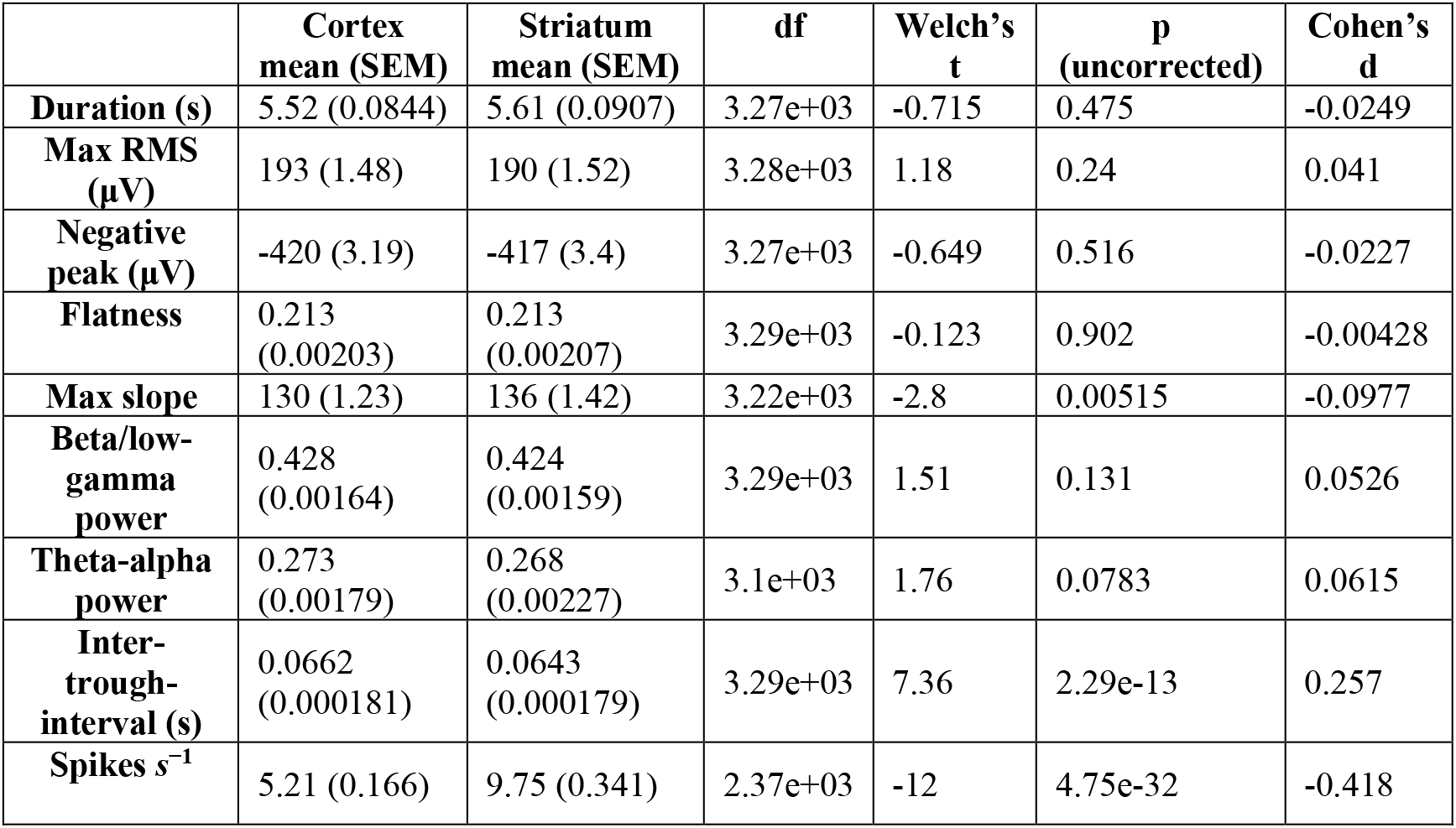
Statistics for cortical versus striatal NGB events.

|  | **Cortex**<br>**mean (SEM)** | **Striatum**<br>**mean (SEM)** | **df** | **Welch's**<br>**t** | **p**<br>**(uncorrected)** | **Cohen's**<br>**d** |
| --- | --- | --- | --- | --- | --- | --- |
| **Duration (s)** | 5.52 (0.0844) | 5.61 (0.0907) | 3.27e+03 | -0.715 | 0.475 | -0.0249 |
| **Max RMS**<br>**(μV)** | 193 (1.48) | 190 (1.52) | 3.28e+03 | 1.18 | 0.24 | 0.041 |
| **Negative**<br>**peak (μV)** | -420 (3.19) | -417 (3.4) | 3.27e+03 | -0.649 | 0.516 | -0.0227 |
| **Flatness** | 0.213<br>(0.00203) | 0.213<br>(0.00207) | 3.29e+03 | -0.123 | 0.902 | -0.00428 |
| **Max slope** | 130 (1.23) | 136 (1.42) | 3.22e+03 | -2.8 | 0.00515 | -0.0977 |
| **Beta/low-**<br>**gamma**<br>**power** | 0.428<br>(0.00164) | 0.424<br>(0.00159) | 3.29e+03 | 1.51 | 0.131 | 0.0526 |
| **Theta-alpha**<br>**power** | 0.273<br>(0.00179) | 0.268<br>(0.00227) | 3.1e+03 | 1.76 | 0.0783 | 0.0615 |
| **Inter-**<br>**trough-**<br>**interval (s)** | 0.0662<br>(0.000181) | 0.0643<br>(0.000179) | 3.29e+03 | 7.36 | 2.29e-13 | 0.257 |
| **Spikes $s^{-1}$** | 5.21 (0.166) | 9.75 (0.341) | 2.37e+03 | -12 | 4.75e-32 | -0.418 |

**Supplemental Table 10:** Statistics for cortical versus thalamic NGB events.

|  | **Cortical**<br>**mean (SEM)** | **Thalamus**<br>**mean (SEM)** | **df** | **Welch's**<br>**t** | **p**<br>**(uncorrected)** | **Cohen's d** |
| --- | --- | --- | --- | --- | --- | --- |
| **Duration**<br>**(s)** | 5.52 (0.0844) | 4.85 (0.0835) | 2.96e+03 | 5.66 | 1.65e-08 | 0.207 |
| **Maximum**<br>**RMS (μV)** | 193 (1.48) | 165 (1.12) | 2.91e+03 | 14.8 | 5.27e-48 | 0.535 |
| **Negative**<br>**peak (μV)** | -420 (3.19) | -406 (3.7) | 2.81e+03 | -2.84 | 0.00458 | -0.105 |
| **Flatness** | 0.213<br>(0.00203) | 0.224<br>(0.00234) | 2.81e+03 | -3.44 | 0.000587 | -0.127 |
| **Maximum**<br>**slope** | 130 (1.23) | 131 (1.3) | 2.91e+03 | -0.132 | 0.895 | -0.00485 |
| **Relative**<br>**beta-low**<br>**gamma**<br>**power** | 0.428<br>(0.00164) | 0.396<br>(0.00194) | 2.78e+03 | 12.5 | 1.03e-34 | 0.46 |
| **Relative**<br>**theta-**<br>**alpha**<br>**power** | 0.273<br>(0.00179) | 0.259<br>(0.0023) | 2.65e+03 | 5.09 | 3.82e-07 | 0.189 |
| **Inter-**<br>**trough-**<br>**interval**<br>**(s)** | 0.0662<br>(0.000181) | 0.0591<br>(0.000169) | 2.98e+03 | 28.7 | 5.16e-160 | 1.05 |
| **Spikes s<sup>-1</sup>** | 5.21 (0.166) | 4.86 (0.146) | 2.99e+03 | 1.56 | 0.12 | 0.0566 |

**Supplemental Table 11:** Statistics for cortical versus thalamic SB events.

|  | **Cortical mean**<br>**(SEM)** | **Thalamus**<br>**mean**<br>**(SEM)** | **df** | **Welch's**<br>**t** | **p**<br>**(uncorrected)** | **Cohen's**<br>**d** |
| --- | --- | --- | --- | --- | --- | --- |
| **Duration**<br>**(s)** | 1.44 (0.0192) | 1.29<br>(0.0233) | 3.03e+03 | 4.72 | 2.47e-06 | 0.161 |
| **Maximum**<br>**RMS (μV)** | 56.1 (0.628) | 50.4 (0.71) | 3.18e+03 | 6.02 | 1.96e-09 | 0.203 |
| **Negative**<br>**peak (μV)** | -126 (1.33) | -115 (1.52) | 3.15e+03 | -5.14 | 2.92e-07 | -0.174 |
| **Flatness** | 0.538<br>(0.00398) | 0.612<br>(0.00474) | 3.08e+03 | -11.9 | 4.32e-32 | -0.405 |
| **Maximum**<br>**slope** | 42.8 (0.377) | 44.7 (0.504) | 2.81e+03 | -3.04 | 0.00242 | -0.104 |
| **Relative**<br>**beta-low**<br>**gamma**<br>**power** | 0.324<br>(0.00181) | 0.315<br>(0.00227) | 2.95e+03 | 3.14 | 0.00171 | 0.107 |
| **Relative**<br>**theta-**<br>**alpha**<br>**power** | 0.268<br>(0.00188) | 0.24<br>(0.00236) | 2.96e+03 | 9.27 | 3.49e-20 | 0.317 |
| **Inter-**<br>**trough-**<br>**interval (s)** | 0.0551<br>(0.000177) | 0.0505<br>(0.000183) | 3.37e+03 | 18.1 | 5.9e-70 | 0.604 |
| **Spikes $s^{-1}$** | 1.35 (0.0508) | 6.51 (0.243) | 1.51e+03 | -20.8 | 7.98e-85 | -0.782 |

**Supplemental Table 12:** Statistics for cortical versus striatal SB events.

|  | **Cortical**<br>**mean (SEM)** | **Striatum**<br>**mean (SEM)** | **df** | **Welch's**<br>**t** | **p**<br>**(uncorrected)** | **Cohen's d** |
| --- | --- | --- | --- | --- | --- | --- |
| **Duration**<br>**(s)** | 1.44 (0.0192) | 1.31 (0.0189) | 4.48e+03 | 4.47 | 8.03e-06 | 0.134 |
| **Maximum**<br>**RMS (μV)** | 56.1 (0.628) | 49.3 (0.56) | 4.42e+03 | 8.08 | 7.97e-16 | 0.242 |
| **Negative**<br>**peak (μV)** | -126 (1.33) | -112 (1.21) | 4.44e+03 | -7.44 | 1.19e-13 | -0.222 |
| **Flatness** | 0.538<br>(0.00398) | 0.576<br>(0.00401) | 4.48e+03 | -6.69 | 2.52e-11 | -0.2 |
| **Maximum**<br>**slope** | 42.8 (0.377) | 41.8 (0.361) | 4.47e+03 | 1.89 | 0.0589 | 0.0565 |
| **Relative**<br>**beta-low**<br>**gamma**<br>**power** | 0.324<br>(0.00181) | 0.306<br>(0.00171) | 4.46e+03 | 7.14 | 1.05e-12 | 0.214 |
| **Relative**<br>**theta-**<br>**alpha**<br>**power** | 0.268<br>(0.00188) | 0.264<br>(0.00193) | 4.47e+03 | 1.53 | 0.125 | 0.0458 |
| **Inter-**<br>**trough-**<br>**interval**<br>**(s)** | 0.0551<br>(0.000177) | 0.0527<br>(0.000162) | 4.44e+03 | 10.2 | 4.32e-24 | 0.304 |
| **Spikes $s^{-1}$** | 1.35 (0.0508) | 2.51 (0.116) | 3.07e+03 | -9.19 | 6.83e-20 | -0.275 |

